# Theoretical framework and experimental demonstration of sustainable in vitro regeneration of major translation factors EF-Tu and IF3

**DOI:** 10.64898/2026.06.14.732184

**Authors:** Kentaro Shoji, Katsumi Hagino, Norikazu Ichihashi

## Abstract

The development of molecular systems capable of self-regeneration, much like living organisms, is a major goal in synthetic biology. Previous efforts to expand the repertoire of regenerating proteins have relied on empirical optimization without a theoretical foundation, fundamentally limiting the scalability of this approach. Here, we developed the first theoretical framework, based on measurable parameters of translational proteins, to rationally predict the dilution rate that enables sustainable regeneration and the steady-state translation level. Using this framework, we successfully demonstrated sustainable regeneration of EF-Tu, the most abundant translation factor, for up to 15 rounds of serial dilution. We further extended this framework to the co-regeneration of multiple translation factors and demonstrated sustainable co-regeneration of EF-Tu and IF3, another major essential translation protein, for up to 20 rounds. These results establish a rational methodology for systematically expanding the number of regenerating proteins, providing a clear path toward the realization of a fully self-regenerating system.

## Introduction

Self-regeneration is an essential property of living systems. The construction of self-regenerating artificial cells is one of the major goals in synthetic biology^1–9^. Although many biological functions required for self-regeneration have been reconstituted in vitro, including DNA replication^10–17^, transcription-translation (TX-TL)^18^, metabolism^19–23^, and membrane synthesis^10,24–26^, a fully self-regenerating system capable of continuously producing all of its own components has yet to be achieved.

To realize the self-regenerating ability in vitro, all genetically encoded macromolecules (DNA, RNA, and proteins) must be synthesized by the system itself through DNA replication, transcription, and translation. The PURE system, a reconstituted cell-free transcription-translation (TX-TL) system, is an ideal platform for this purpose owing to its minimal, defined, and adjustable composition^18^. The PURE system contains all components required for TX-TL, including ribosomes, tRNAs, T7 RNA polymerase (T7 RNAP), translation factors (TFs) such as aminoacyl-tRNA synthetases (aaRSs), translational initiation, elongation, and releasing proteins. Therefore, regeneration of all PURE system components from DNA within the PURE system would represent the first step toward constructing a fully self-regenerating system^27,28^.

Previous studies on the self-regeneration of the PURE system can be broadly categorized into three types. The first type focused on the expressed level of TFs from DNA without measuring their activities. For example, Doerr et al. reported simultaneous synthesis of 32 TFs in the PURE system from a large DNA^29^, and Libicher et al. reported expression of at least 30 TFs from a DNA set coupled with DNA replication^30^. The second type measured the translation activities of newly synthesized TFs produced from DNA either *in situ* within the PURE system^31,32^ or in a separate reaction^33–35^. For example, Schwarz-Schilling et al. recently detected GFP expression driven by in-situ-expressed 30 TFs^32^ or by a ribosomal subunit assembled in situ^36^. The primary purpose of this approach is simply to detect the activity of the newly synthesized factors without regard for quantitative activity levels. However, to achieve sustainable regeneration as living systems do, the regenerated TFs must maintain the steady-state activity. The third type aims to maintain translational activity to a steady-state level by synthesizing the lacking TFs. For example, Maerkl’s group reported sustainable regeneration of seven aaRSs for 22 h using microfluidics^37^. We also reported the regeneration of all 20 aaRSs for 20 rounds of 3-fold serial dilution^38^. However, sustainable regeneration has so far remained limited to a minor fraction of non-ribosomal TFs in the PURE system (aaRSs account for only 2.5% of the total TF composition; Fig. 2a). The next challenge is the regeneration of major TFs, such as EF-Tu and other elongation and initiation factors.

To achieve sustainable regeneration of major TFs, we argue that a more systematic approach is needed. Previous attempts at sustainable regeneration have largely relied on empirical optimization of reaction conditions in an ad hoc manner, at least for our case. This conventional approach makes the expansion to major TFs difficult because their regeneration requires higher translation activity, making it more difficult to identify conditions that support regeneration. Therefore, a theoretical framework that can rationally predict the conditions required for successful regeneration is essential. Such a framework should be capable of predicting the success or failure of sustainable regeneration from quantitative parameters (e.g., expression levels, activity per protein) of each TF without requiring actual regeneration experiments. This approach will enable research to proceed in a predictable and rational manner, rather than relying on ad hoc, trial-and-error optimization.

Here, we developed a theoretical framework to assess the feasibility of sustainable self-regeneration of TFs using a serial dilution method. In this framework, the feasibility of sustainable regeneration is determined based on an expression curve that describes the relationship between the target protein concentrations before and after a single round of the TX-TL reaction. If the expression curve intersects a dilution line, the TF is sustainably regenerated at the concentration of the intersection point. This framework was initially applied to EF-Tu, the most abundant TF in our PURE system composition, and then extended to co-regeneration with IF3, the second most abundant essential TF. In each case, we predicted the range of dilution rate that allows sustainable regeneration, and then experimentally demonstrated the sustainable regeneration for 15–20 rounds of serial dilution. Using this framework, we further discussed requirements for realizing sustainable regeneration of all TFs.

## Results

### Theoretical framework for sustainable regeneration

To formulate a theoretical framework for sustainable regeneration, we first needed to determine the method for supplying substrates (e.g., amino acids and NTPs) and non-regenerating TFs, because without their replenishment, protein synthesis eventually ceases and regeneration cannot be sustained. In previous studies, substrates and TFs have been supplied by two methods: the serial dilution method^31,38^ or the continuous method using microfluidics^37^. Here, we focused on the serial dilution method owing to its technical simplicity (Fig. 1a). In this study, we defined “sustainable regeneration” as the synthesis of target TFs at a level sufficient to maintain a steady-state translational activity throughout repeated serial dilution of the PURE system lacking the target TFs. We next formulated the requirements for achieving sustainable regeneration of a TF in the PURE system. To achieve sustainable regeneration in a serial dilution format, target proteins must be synthesized in sufficient quantities in a single TX-TL reaction to compensate for the dilution losses. More specifically, the total active TF concentration after the single TX-TL reaction must be at least D-fold (D being the dilution factor, which must be >1) greater than the initial TF concentration before the reaction. This requirement can be represented as a graph of the relationship between the TF concentration before and after a single round of the TX-TL reaction, referred to as an “expression curve” in this study (blue line in Fig. 1b). The expression curve typically exhibits a concave shape because TFs become sufficient for translation at higher concentrations. If this expression curve intersects a dilution line (y = Dx), the TF is sustainably regenerated by D-fold serial dilution, and the intersection point represents the steady-state TF concentrations under sustainable regeneration.

**Figure 1.**
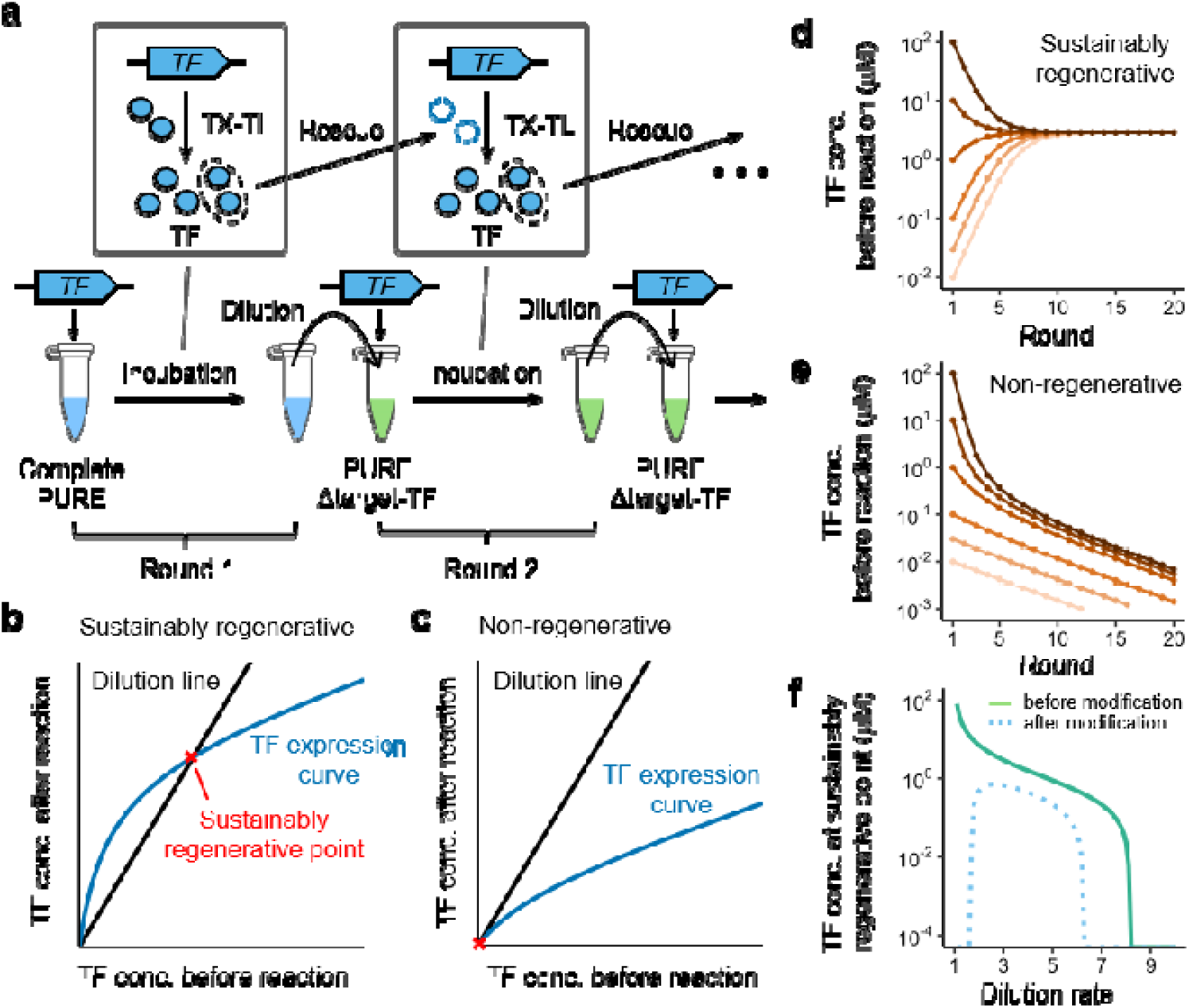
Theoretical framework for sustainable regeneration of a TF. (a) Schematic of the serial dilution method for sustainable regeneration. First, the target TF gene is expressed in the PURE system containing all TFs (Complete PURE) at round 1. A fraction of the reaction mixture is then diluted with the PURE system lacking the target TF (PURE Δtarget-TF) from round 2 onward. This cycle of reaction and dilution was repeated. If a sufficient amount of functional TF is synthesized, translational activity is maintained in subsequent rounds, allowing continued TF synthesis. (b and c) Conceptual framework for assessing the feasibility of sustainable regeneration. The blue TF expression curve represents the TF concentration after the single round of TX-TL reaction as a function of the TF concentration before the reaction. The black dilution line has a slope corresponding to the dilution rate. The target TF can be sustainably regenerated when these lines intersect (b), but not when they do not intersect (c). (d and e) Simulated TF concentration trajectories during serial dilution with varying initial TF concentrations. The TF expression curves obtained from later experiments (Fig. 6) were used for simulations under conditions where the expression curve intersects the dilution line (d, 3-fold dilution) or does not intersect (e, 10-fold dilution). (f) Relationship between dilution rates and the steady-state TF concentrations at the sustainably regenerative points. Solid and dotted lines represent the relationships without and with considering the accumulation of inhibitory factors, respectively.

This interpretation can be explained as follows. When an intersection exists, and the expression curve is higher than the dilution line (i.e., in the left of the intersection in Fig. 1b), TF production is higher than the dilution loss, leading to TF accumulation toward the intersection. In contrast, when the expression curve is lower than the dilution line (i.e., in the right of the intersection), TF production is lower than the dilution loss, leading to TF depletion toward the intersection point. As a result, the TF concentration stabilizes at the intersection, where synthesis and dilution loss are balanced (see Fig. S1 for a more detailed explanation). However, if the dilution rate is too large and no intersection exists, the TF concentration gradually decreases and eventually reaches zero (Fig. 1c). To confirm this prediction, we performed computer simulations using the expression curve of EF-Tu obtained experimentally (see below). When an intersection existed, the TF concentrations converged to the steady-state concentration defined by the intersection point, regardless of the initial TF concentration (Fig. 1d), while they continuously decreased when no intersection existed (Fig. 1e).

Given an expression curve, we can further predict the relationship between the dilution rate and the steady-state (i.e., sustainably regenerative) TF concentration. Using the expression curve of EF-Tu obtained later as an example, a steady-state TF concentration is achieved when the dilution rate is less than approximately eight, with the concentration increasing as the dilution rate decreases (Fig. 1f, green solid line). As discussed later, this relationship is modified by the accumulation of inhibitory factors during serial dilution (Fig. 1f, blue dotted line), which further narrows the range of dilution rates that allow sustainable regeneration to a limited window.

In summary, according to this theoretical framework, the feasibility of sustainable regeneration is determined by whether the TF expression curve intersects a dilution line. Because the dilution line is arbitrarily determined by the dilution rate, the feasibility of sustainable regeneration can be assessed once the TF expression curve has been obtained. In the following sections, we first determined the target TFs and then performed a series of characterizations to obtain the precise expression curve.

### Selection of target TFs to be regenerated

Fig. 2a shows the composition of TFs in our customized PURE system (ver. 2 of Kazuta et al.^39^), which exhibits significantly higher translation activity than the original composition^18^. To prioritize target TFs for regeneration, we first assessed the essentiality of the major TFs (IF1, IF2, IF3, EF-Ts, EF-Tu, EF-G, RF1, RF2, RF3, and RRF) for translation by measuring GFP expression in a customized PURE system lacking each individual TF. Omission of IF2, IF3, EF-G, and EF-Tu significantly decreased GFP expression (Fig. 2b), indicating that these TFs are essential for translation in our PURE system. Based on these results, we selected EF-Tu and IF3, the first and second most abundant essential TFs, as the targets for regeneration in this study. Although the composition of TFs varies among different PURE systems (Fig. S2), EF-Tu and IF3 are consistently major factors. We did not choose IF1 because it was not essential, consistent with previous reports^32,34,40^.

**Figure 2.**
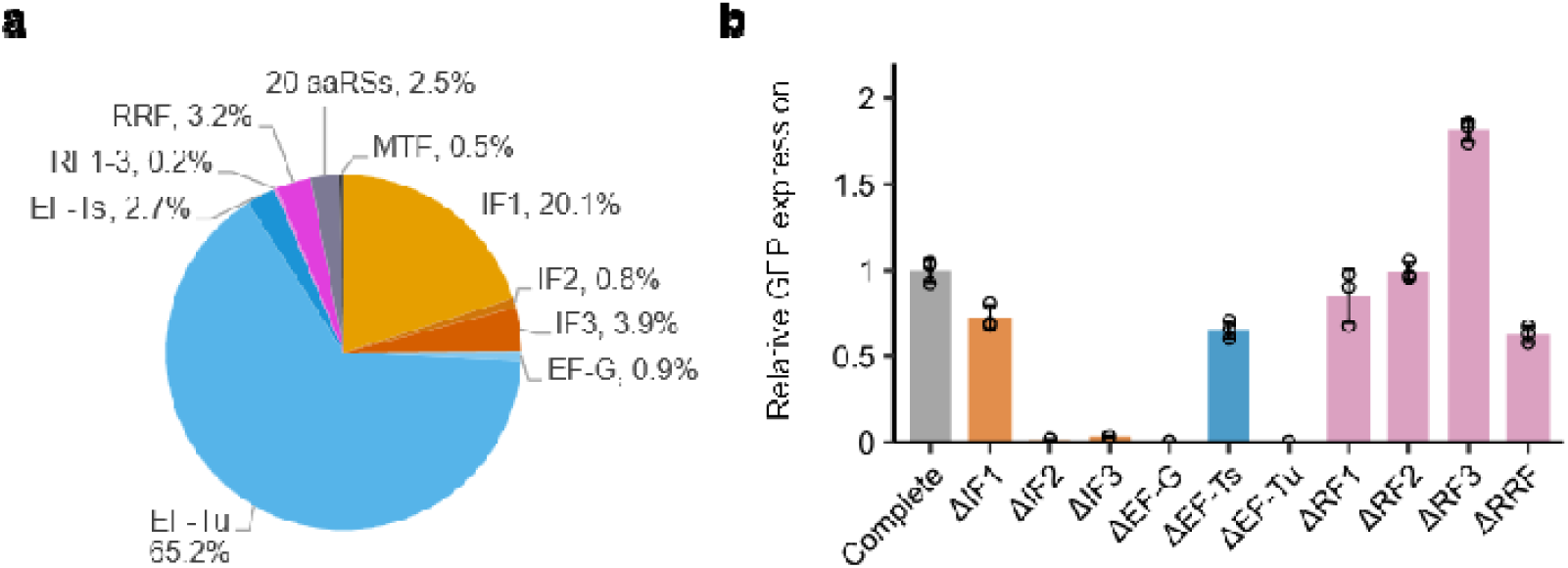
Composition of TFs in our PURE system and their essentiality. (a) TF composition of the PURE system used in this study, calculated from molar concentrations. TF compositions of other PURE systems are shown in Fig. S2. (b) Essentiality of TFs for translation. GFP encoding DNA (3 nM) was added to the PURE system lacking each TF (ΔTF) and incubated at 30°C. For ΔRF1 and ΔRF2, GFP encoding DNAs containing stop codons specifically recognized by RF1 or RF2 were used. GFP fluorescence was measured every 15 min. The time-course data of GFP fluorescence are shown in Fig. S3. The increase in GFP fluorescence after 8 h was normalized to the value obtained with the PURE system containing all TFs (Complete PURE). Bars indicate mean values ± standard deviation (SD) from three independent experiments. IF, initiation factor; EF, elongation factor; RF, release factor; RRF, ribosome recycling factor; MTF, methionyl-tRNA formyltransferase.

### Characterization of EF-Tu and IF3 expression and activity

To obtain the precise expression curve, we required three types of quantitative data for each of the two target TFs, EF-Tu and IF3: 1) the expression level of each target TF in the PURE system, 2) the translation activity of the expressed protein, and 3) the effect of each target TF concentration on translation. Furthermore, to design regeneration experiments based on the expression curve, we required two additional types of quantitative data: 4) the effect of the DNA concentration of each target TF on the gene expression, and 5) the effect of the DNA ratio of the two TFs on translation during co-regeneration. We obtained these data as follows.

First, we characterized the expression levels of EF-Tu and IF3 produced from DNA in the PURE system (data type 1). We prepared three variants of each TF: the original tagged version (+His), a histidine-tag-removed version (ΔHis), in which the histidine tag attached to the N- or C-terminus for purification was removed, and a codon optimized version of the latter (ΔHis-opt), generated using an algorithm^41^, with the aim of enhancing the expression levels and functional activity of these TFs (Fig. 3a). These variants were expressed in our customized PURE system and their expression levels were measured using a HiBiT tag attached to the 3′-terminus. No significant differences were observed among the EF-Tu variants, while the IF3 (ΔHis) variant exhibited slightly higher expression levels (Fig. 3b).

**Figure 3.**
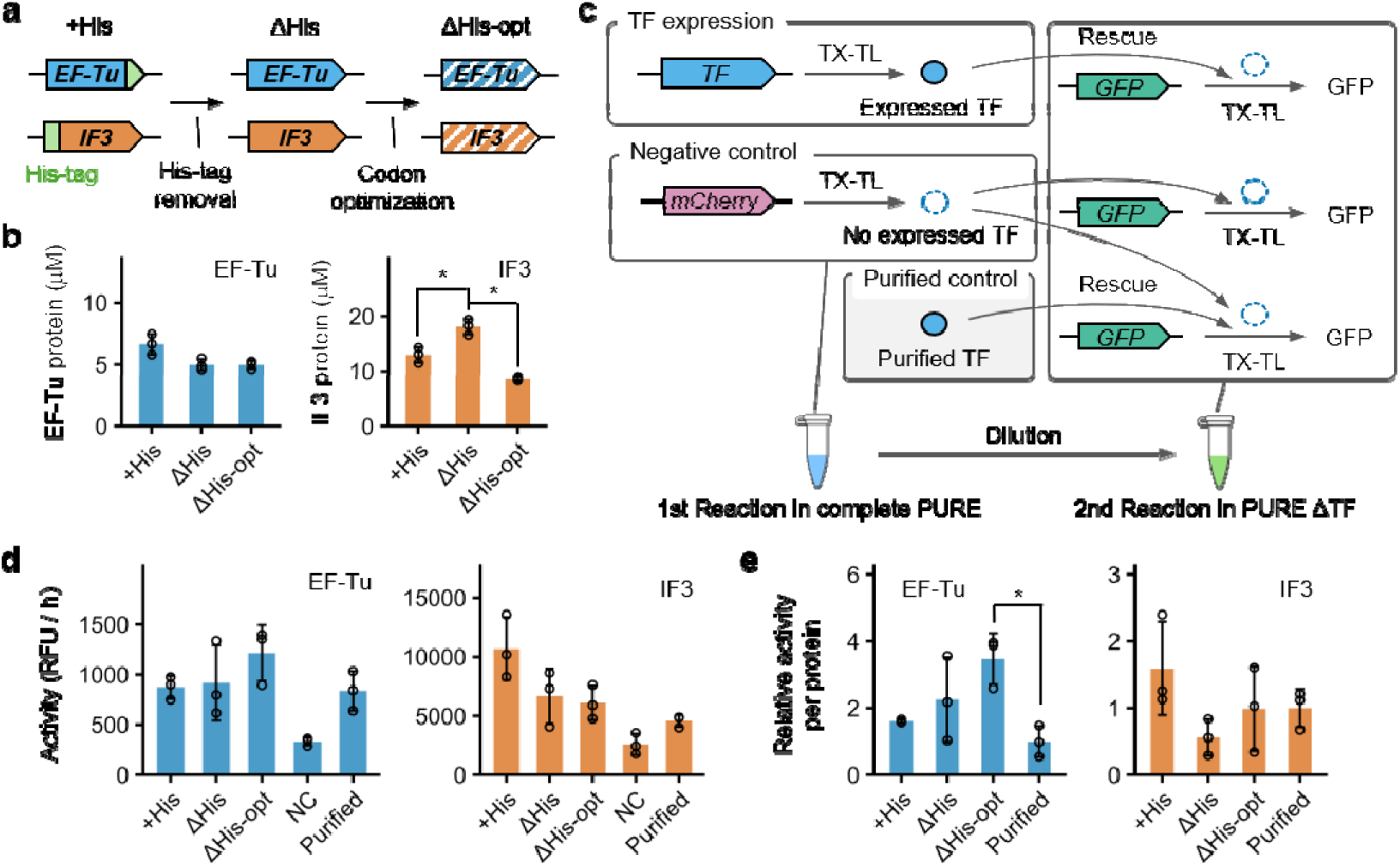
Expression levels and the activities of expressed EF-Tu and IF3 variants. (a) Schematic of EF-Tu and IF3 variants. The three variants include the conventional construct containing a histidine tag (+His), a construct lacking the His-tag (ΔHis), and a construct further modified by codon optimization (ΔHis-opt). (b) Expression levels of each variant. Template DNAs encoding the modified variants with a C-terminal HiBiT tag (4 nM) were added to the PURE system and incubated at 30°C for 8 h. Expression levels were quantified using the HiBiT assay. (c) Schematic of the TF activity assay for TFs expressed in the PURE system. The assay consisted of two steps. First, TF encoding DNA (4 nM) was expressed in the complete PURE system (1^st^ reaction); mCherry encoding DNA was expressed as a negative control (NC). Second, dilution series of the 1^st^ reaction mixture were prepared and added to PURE systems lacking the target TF and containing GFP encoding DNA (3 nM), followed by GFP expression (2^nd^ reaction). Both reactions were performed at 30°C for 8 h. For activity measurements of purified TFs, purified TFs were added to the NC 1^st^ reaction mixture. (d) Activity measurements of TFs expressed from each variant and of purified TFs. GFP fluorescence was measured every 10 min (time-course data are shown in Fig. S4). GFP synthesis rates were plotted against the dilution ratios of the 1^st^ reaction mixtures (shown in Fig. S5), and the slopes calculated by linear regression were defined as TF activity. (e) Activity per protein of each variant. TF activities in (d) were subtracted by the NC value and divided by the corresponding TF expression levels in (b). Activities of purified TFs were divided by the concentrations added in the 2^nd^ reaction. The calculated values were normalized to those of purified TFs. (b, d, e) Bars indicate mean values ± SD from three independent experiments. Statistical significance was determined using Welch’s t-test with Bonferroni correction for multiple comparisons. \**p* < 0.05.

Second, we characterized the translational activity of the expressed protein (data type 2). We assessed the functional activity of PURE-expressed EF-Tu and IF3 relative to that of purified proteins using the activity assay illustrated in Fig 3c. In this assay, each TF variant was expressed in a complete or TF-reduced PURE system, with mCherry DNA added as a negative control to impose a translational burden (1^st^ reaction). The resulting mixtures were then diluted at various ratios and added to a PURE system lacking the corresponding TF, into which GFP DNA was introduced to evaluate translational activity via GFP fluorescence (2^nd^ reaction). For comparison, the activities of the purified proteins were measured by adding them to the 2^nd^ reaction mixture with the 1^st^ reaction product of mCherry. The GFP synthesis rates during the 2^nd^ reactions were then measured during the linear phases (Fig. S4) and plotted against the dilution ratios of the 1^st^ reaction mixtures (Fig. S5). The resulting slopes were used to evaluate the activity of each TF variant (Fig. 3d). EF-Tu showed the highest activity with the ΔHis-opt variant, while IF3 showed the highest activity with the +His variant, although the differences were not statistically significant. We then calculated the activity per protein by dividing the activity values in Fig. 3d by the corresponding expression levels in Fig. 3b after subtracting the negative control value (Fig. 3e). EF-Tu ΔHis-opt showed the highest activity per protein, which was 3.5-fold higher than that of the purified EF-Tu. In contrast, IF3 showed the highest activity per protein with the +His variant, at 1.6-fold higher than that of the purified IF3, although this difference was not statistically significant. Based on these results, we selected the EF-Tu (ΔHis-opt) and the IF3 (+His) for subsequent experiments.

### Effect of protein and DNA concentrations of EF-Tu and IF3 on translation

Third, we measured the effect of a target TF concentration on translation (data type 3). These data were obtained by measuring GFP expression using purified EF-Tu (+His) and IF3 (+His) at various concentrations (Fig. 4a). The GFP fluorescence values were normalized to the values obtained at the original concentrations (EF-Tu: 80 μM; IF3: 5 μM). Each curve was fitted to a Michaelis–Menten-type saturation function to quantify the characteristic parameters, *K*_1/2_ and *A*_max_ (Fig. 4a). The significantly larger *K*_1/2_ of EF-Tu reflects a greater requirement for EF-Tu in translation, consistent with its high concentration in our PURE system.

**Figure 4.**
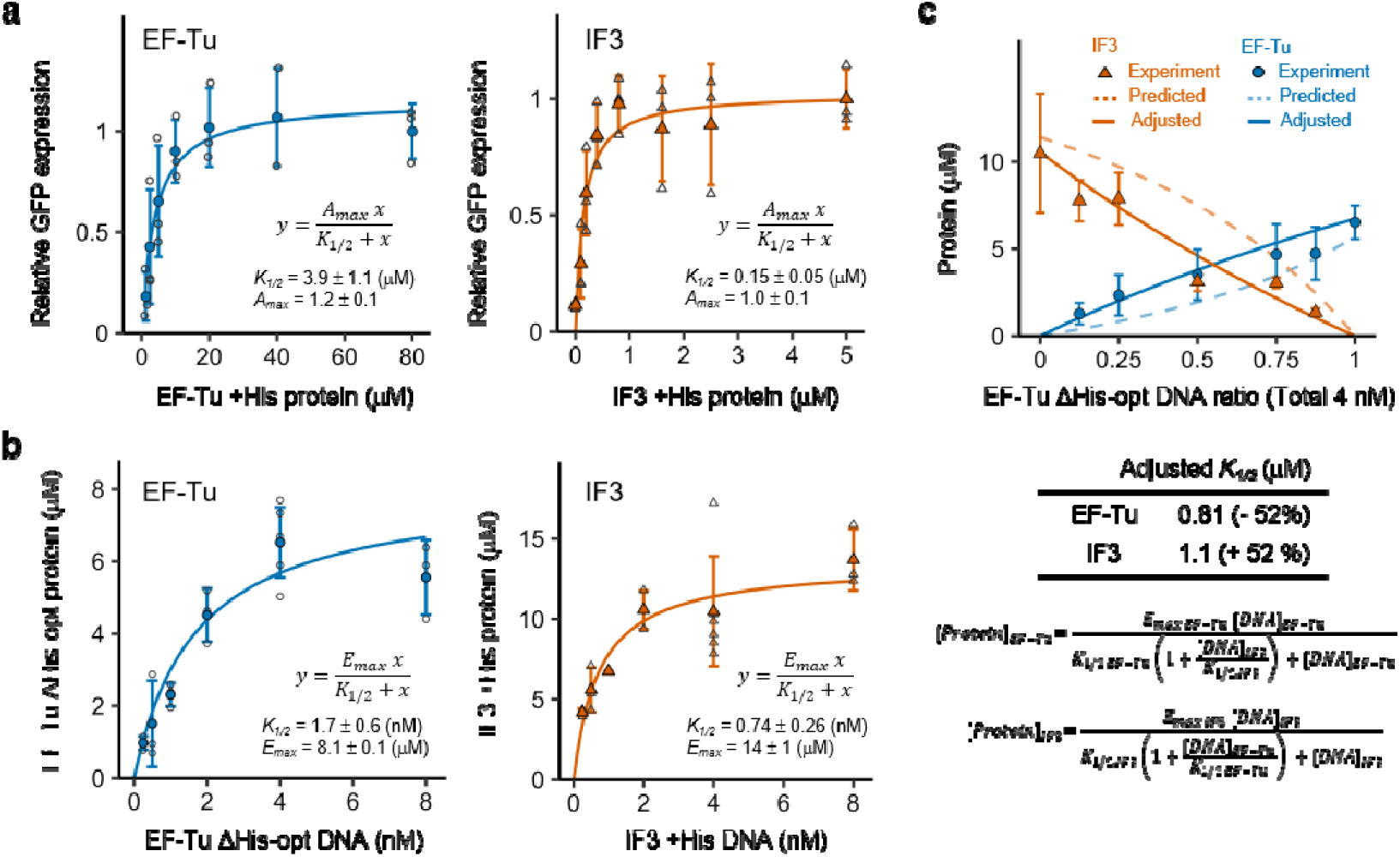
Effect of TF protein or DNA concentrations on expression level. (a) Dependence of translational activity on each purified TF concentration. GFP encoding DNA (3 nM) and varying concentrations of each purified TF were added to the PURE systems lacking the corresponding TF and incubated at 30°C. After 6–8 h, the increases in GFP fluorescence were normalized to the values obtained at the TF concentrations in Complete PURE (EF-Tu, 80 μM; IF3, 5 μM). Data points represent mean values ± SD from two or three independent experiments. Curves fitted to a Michaelis-Menten-type saturation function are shown. (b) Dependence of TF expression levels on DNA concentration. The TF encoding DNAs bearing HiBiT tags were added to the PURE system at varying concentrations and incubated at 30°C for 8 h. Expression levels were quantified using the HiBiT assay. Data points represent mean values ± SD from three to six independent experiments. Curves fitted to a Michaelis-Menten-type saturation function are shown. (c) Co-expression experiment of EF-Tu and IF3. EF-Tu and IF3 encoding DNAs were mixed at various ratios (a total of 4 nM) and expressed in the PURE system at 30°C for 8 h. The expression levels of each TF were quantified using a HiBiT-tag attached to one of the TF encoding DNAs. Dotted lines show predictions generated using a competitive inhibition model (equation is shown at the bottom) with parameters from Fig. 4b. Solid lines show predictions after adjusting *K_1/2_*. Data points represent mean values ± SD from three independent experiments.

Fourth, we measured the effect of each TF encoding DNA concentration on the expressed level (data type 4). These data were obtained by measuring the expression level of each TF using a HiBiT tag at various DNA concentrations (Fig. 4b). Each curve was fitted to a Michaelis-Menten-type saturation function to quantify the characteristic parameters, *K*_1/2_ and *E*_max_ (Fig. 4b). The larger *K*_1/2_ for EF-Tu indicates that more DNA is required to achieve the maximum expression, while the smaller *E*_max_ for EF-Tu indicates a lower maximum expression level. These data were used to determine the DNA concentration required for the regeneration experiments described later.

Fifth, we measured the effect of the DNA ratio of the two TFs on the expression level of each protein during co-expression (data type 5). The two DNA templates were mixed at various ratios (total 4 nM), and the expression level of each was quantified using a HiBiT tag (Fig. 4c). In theory, if the two DNAs compete for the limited transcriptional/translational resources, the expression levels can be calculated using the formula shown in Fig. 4c (see Methods for derivation) using the parameters *K_1/2_*and *E_max_* obtained in Fig. 4b; however, the calculated curves (dotted lines in Fig. 4c) were slightly lower than the actual EF-Tu expression levels and higher than the actual IF3 expression levels, probably due to unknown interactions between EF-Tu and IF3 expressions during co-expression. We therefore adjusted the *K_1/2_* values of EF-Tu and IF3 proportionally to best fit the experimental data (Fig. 4c, solid lines). These adjusted parameters were used to determine the DNA concentrations required for the co-regeneration experiments described later.

### Characterization of the decreasing translational activity during serial dilution

During optimization of the serial dilution method, we noticed that the translation activity gradually decreased when using a higher concentration of T7 RNA polymerase (T7 RNAP), which catalyzes transcription in the PURE system. Since this decrease could influence the feasibility of sustainable regeneration, we sought to quantitatively evaluate it before conducting the regeneration experiments. We performed a serial dilution experiment in which an aliquot of the complete PURE system containing EF-Tu and a reporter luciferase encoding DNAs was serially diluted with a fresh complete PURE system containing the same DNA set after incubation at 30°C for 8 h (Fig. 5a). Here, the EF-Tu encoding DNA was included to mimic the translational burden of TF regeneration. We conducted the experiment for three rounds of 3-fold serial dilution at various T7 RNAP concentrations (6.25, 12.5, 25, and 50 nM) and quantified luciferase activity as a measure of translational activity at the end of each round (Fig. 5b).

**Figure 5.**
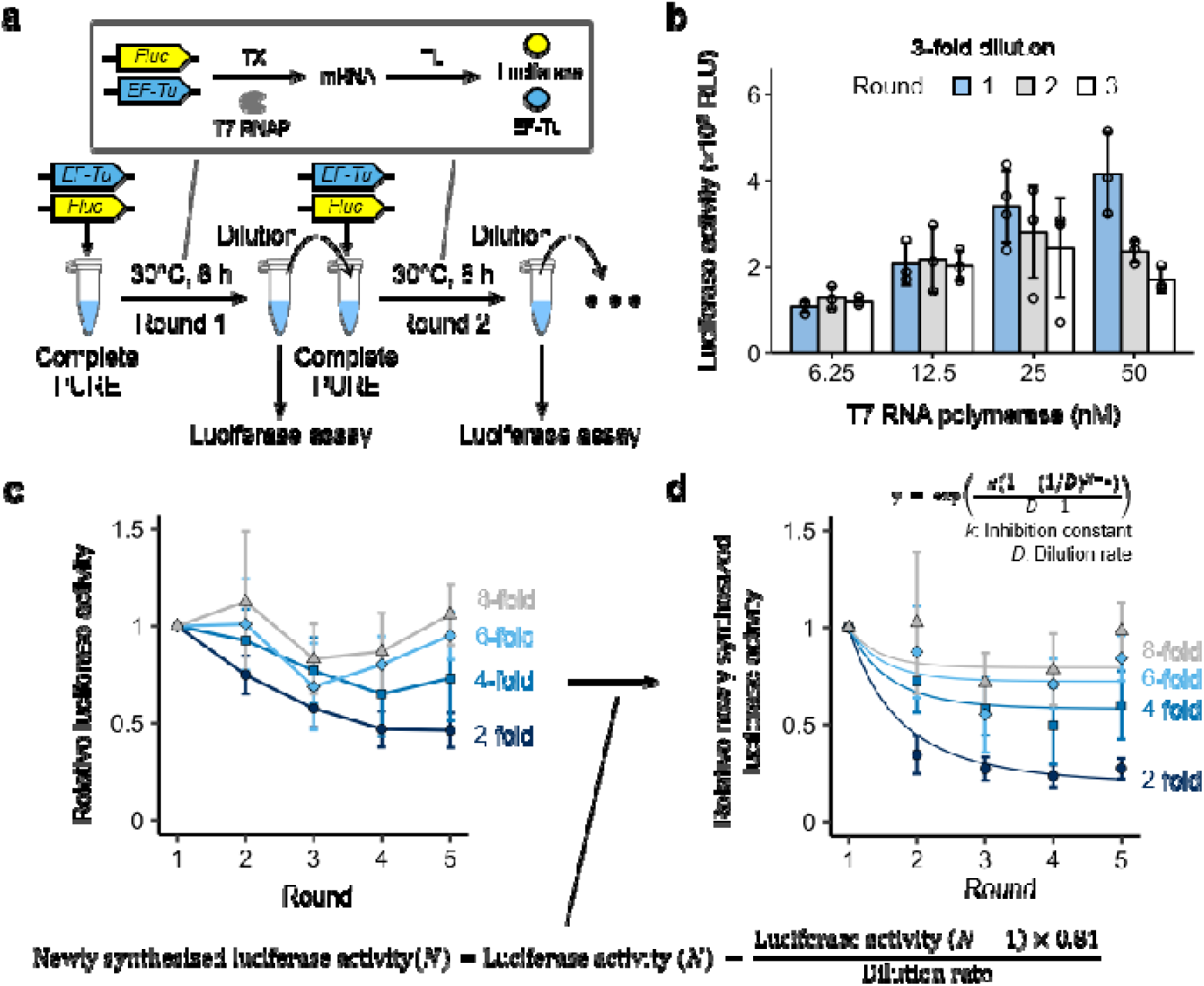
Effect of T7 RNAP concentration and dilution rate on steady-state translation activity during serial dilution. (a) Schematic of the serial dilution experiment. EF-Tu encoding DNA (4 nM) and firefly luciferase (Fluc) encoding DNA (0.01 nM) were added to the Complete PURE system containing all TFs and incubated at 30°C for 8 h. The reaction mixture was then diluted at a defined dilution rate with fresh Complete PURE system. Luciferase activity was measured after each round. (b) Effect of T7 RNAP concentration on luciferase activity during serial dilution. Bars indicate mean values ± SD from three or four independent experiments. (c) Effect of dilution rate on luciferase activity during serial dilution. Data points represent mean values ± SD from two or three independent experiments. (d) Newly synthesized luciferase activity at each round for different dilution rates. For each round, the luciferase activity carried over from the previous round was subtracted from the total luciferase activity to estimate the activity of newly synthesized luciferase. The resulting values were fitted to a model assuming the accumulation of inhibitory byproducts during serial dilution (solid lines).

Translational activity increased with T7 RNAP concentration in round 1, while it declined as the rounds proceeded, especially at higher T7 RNAP concentrations (25 and 50 nM) (Fig. 5b). A similar result was obtained with 2-fold dilutions (Fig. S6). The highest translational activity in round 3 was obtained with 25 nM T7 RNAP, which is lower than the optimal concentration at round 1 (50 nM), indicating that the optimum T7 RNAP concentration is lower at the steady-state of the serial dilution process. Therefore, we selected 25 nM T7 RNAP for all subsequent regeneration experiments.

The gradual decrease in translational activity during serial dilution may be caused by the accumulation of inhibitory factors, likely RNA-related, because the decrease correlates with the T7 RNAP concentration. If this is the case, the dilution rate should affect the degree of decline in translational activity. To test this, we performed the same serial dilution experiments using a fixed T7 RNAP concentration (25 nM) at 2-, 3-, 4-, and 8-fold dilutions for five rounds. The translational activity gradually decreased over successive rounds, and the decrease was more pronounced at lower dilution rates (Fig. 5c). At 2- and 4-fold dilution, the luciferase activity decreased to 46% and 72% of the round 1 level by round 5, respectively. These substantial decreases in translational activity must therefore be incorporated into our theoretical framework.

From these data, we estimated the steady-state translational activity at each dilution rate. To accurately quantify the newly synthesized luciferase activity in each round, we subtracted the luciferase activity carried over from the previous round after accounting for the inactivation rate during incubation (see Methods for detailed calculations). The calculated translational activities revealed sharp decreases in round 2, followed by approximately constant levels (Fig. 5d). To estimate the steady-state translational activity as a function of the dilution rate, we fitted the data to the equation shown in Fig 5d, which assumes the accumulation of inhibitory byproducts, to obtain the inhibition constant *k* (= 1.61) (lines in Fig. 5d; see Methods).

### Derivation of the expression curve for EF-Tu

Having obtained all required data, we estimated the expression curve to predict the dilution rate that enables sustainable regeneration of EF-Tu. The derivation procedure is shown in Fig. 6. We started from the fitted curve of relative GFP expression as a function of purified EF-Tu +His protein concentration obtained from Fig. 4a (Fig. 6a). First, we converted the x-axis from the purified EF-Tu +His concentration to the EF-Tu ΔHis-opt concentration (Fig. 6b), based on the relative activity per protein of the EF-Tu ΔHis-opt variant obtained in Fig. 3e, which reduced *K_1/2_* to 1/3.4 of its original value. Next, we converted the y-axis from relative GFP expression to the EF-Tu ΔHis-opt expression level (Fig. 6c), using the maximum expression level of EF-Tu ΔHis-opt (8.1 µM) obtained in Fig. 4b. We then converted the y-axis to reflect EF-Tu expression during serial dilution (Fig. 6d), using the steady-state translational activity values at 2-, 3-, 4-, and 8-fold dilutions calculated from the model used in Fig. 5d. Finally, we converted the y-axis to represent the total EF-Tu concentration after the reaction, defined as the sum of the initial EF-Tu concentration and the newly expressed EF-Tu (Fig. 6e), yielding the expression curves describing the EF-Tu ΔHis-opt protein concentration before and after the reaction for each dilution rate.

**Figure 6.**
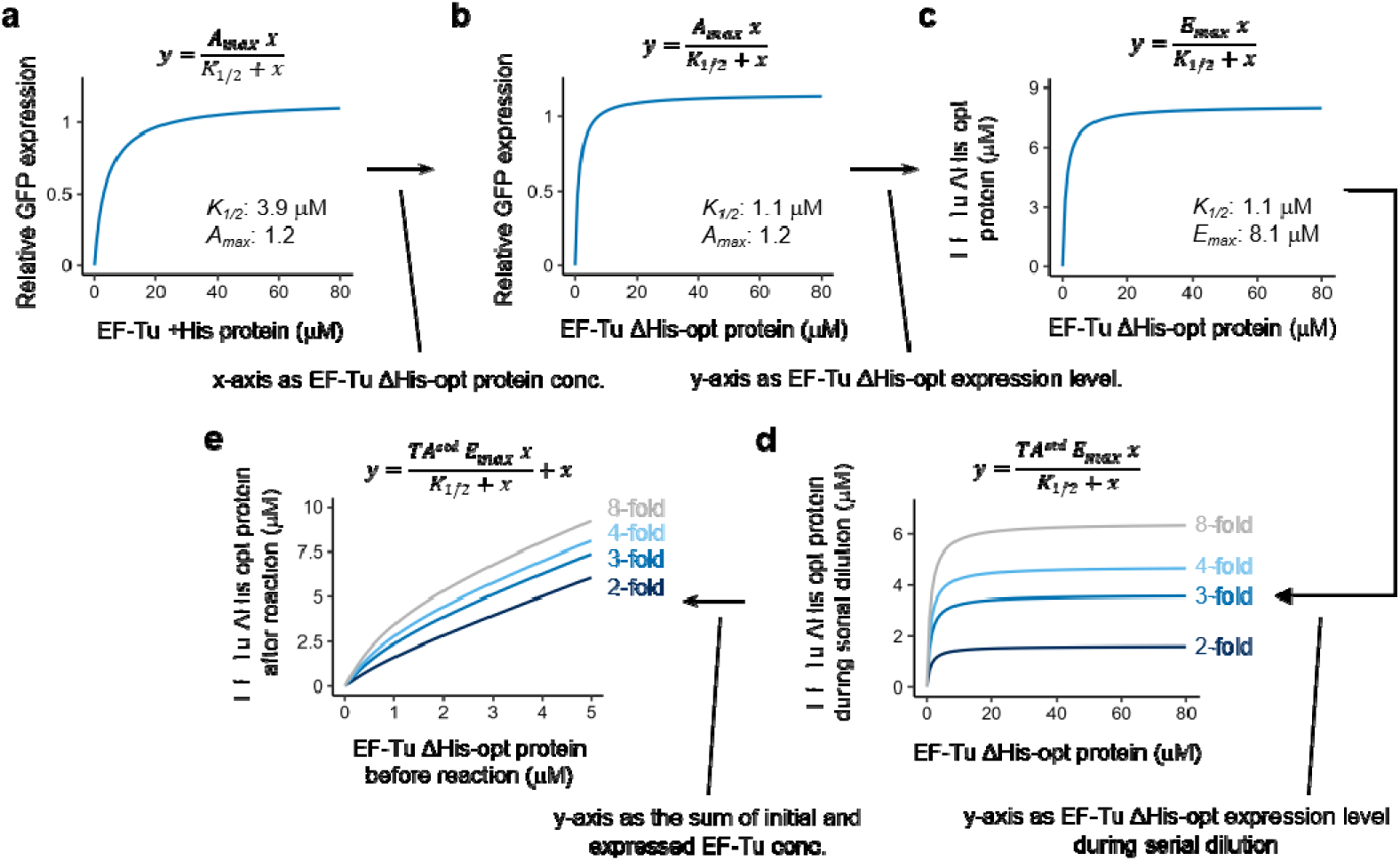
Derivation of the expression curve for EF-Tu. (a) Relative GFP expression as a function of purified EF-Tu +His concentration. The solid curve represents a Michaelis–Menten-type saturation function fitted to the experimental data in Fig. 4a. (b) Relative GFP expression as a function of EF-Tu ΔHis-opt concentration. The x-axis in (a) was converted to the EF-Tu ΔHis-opt concentration by dividing by the relative activity per protein of the ΔHis-opt variant in Fig. 3e. (c) EF-Tu ΔHis-opt expression level as a function of EF-Tu ΔHis-opt concentration. The y-axis in (b) was converted to the EF-Tu ΔHis-opt expression level by using the maximum expression level of EF-Tu ΔHis-opt obtained in Fig. 4b. (d) EF-Tu ΔHis-opt expression during serial dilution as a function of EF-Tu ΔHis-opt concentration at various dilution rates. The y-axis in (c) was multiplied by the steady-state translational activity (TA^std^) at each dilution rate shown in Fig. 5d to estimate the EF-Tu ΔHis-opt expression levels at the steady-state of serial dilution. (e) EF-Tu ΔHis-opt concentration before and after the reaction during serial dilution. The y-axis in (d) was converted to the post-reaction EF-Tu ΔHis-opt concentration by adding the pre-reaction EF-Tu ΔHis-opt concentration.

By overlaying this expression curve with the dilution lines (*y = Dx*; *D* = 2, 3, 4, and 8), we determined the feasibility of sustainable regeneration based on the presence or absence of intersection points (Fig. 7a). At 3-fold dilution, for example, the expression curves intersected the dilution line, predicting that EF-Tu would be stably regenerated at pre-and post-reaction concentrations of 0.67 μM and 2.0 μM, respectively. Similarly, the expression curves intersected the dilution lines at 2- and 4-fold dilutions but not at 8-fold dilution (Fig. S7). The predicted steady-state EF-Tu concentration before the reaction (i.e., the x-value of the intersection point) as a function of dilution rate is shown in Fig. 7b (dotted line). The presence of intersection points in the dilution rate range of 1.7 – 6.2 predicts that EF-Tu can be sustainably regenerated within this range, at the indicated steady-state concentrations.

**Figure 7.**
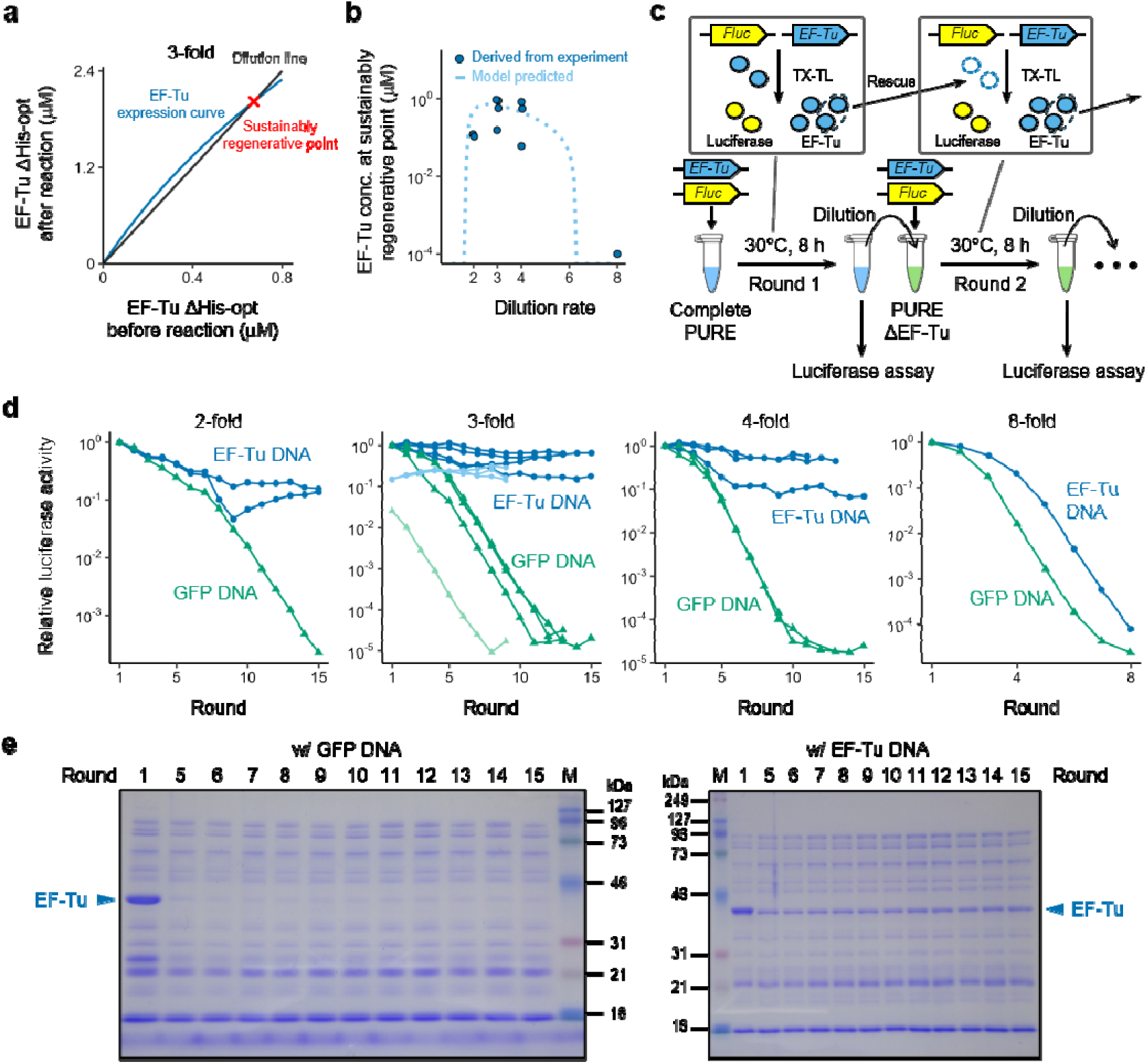
Prediction and experimental verification of sustainable EF-Tu regeneration. (a) Theoretical prediction of EF-Tu regeneration at 3-fold dilution. The black line represents the dilution line (y = 3x), and the blue curve represents the EF-Tu expression curve at 3-fold dilution derived in Fig. 6e. The intersection point is indicated by a red cross. (b) Theoretical predictions and experimentally estimated steady-state EF-Tu concentrations during serial dilution. The dashed line represents the predicted EF-Tu concentration before the reaction at the steady state. Blue circles indicate EF-Tu concentrations derived from the experimentally measured luciferase activities at the end of the serial dilution experiments in Fig. 7d. (c) Schematic of the EF-Tu serial dilution experiment. In the first round, EF-Tu encoding DNA (4 nM) and Fluc encoding DNA (0.01 nM) were expressed in the complete PURE system. In subsequent rounds, the reaction mixtures were diluted at 2-, 3-, 4-, and 8-fold with the PURE system lacking EF-Tu (PURE ΔEF-Tu) and containing the same DNA sets. All reactions were performed at 30°C for 8 h. Luciferase activity was measured after each round to assess translational activity. (d) Trajectories of luciferase activity at 2-, 3-, 4-, and 8-fold dilutions. Luciferase activities were normalized to the round 1 value. The light-colored line for the 3-fold dilution condition represents an experiment in which the initial EF-Tu concentration was reduced to 0.625 μM in round 1. (e) Verification of EF-Tu synthesis by SDS-PAGE. The reaction mixtures before dilution at the indicated rounds were subjected to SDS-PAGE on a 9% gel and stained with Coomassie Brilliant Blue (CBB). The band corresponding to EF-Tu is indicated by an arrowhead. Band intensities were quantified in Fig. S8.

### Regeneration experiments of EF-Tu

To experimentally verify these predictions, we performed serial dilution experiments for up to 15 rounds at 2-, 3-, 4-, and 8-fold dilutions, using EF-Tu encoding DNA (4 nM) and luciferase encoding DNA (0.01 nM). As a negative control, GFP encoding DNA was used instead of EF-Tu encoding DNA. The experimental procedure is shown in Fig. 7c. The first round was carried out in the complete PURE system to initiate EF-Tu expression. In subsequent rounds, the reaction mixture was diluted with the ΔEF-Tu PURE system. As the initial EF-Tu present in the complete PURE system is progressively diluted with each round, translational activity would be lost in the absence of de novo EF-Tu synthesis from the DNA template. Therefore, the maintenance of translational activity serves as an indicator that newly synthesized EF-Tu is functional and capable of supporting its own expression in subsequent rounds.

Consistent with our prediction, luciferase activity was maintained at approximately 5.8% of the initial level for up to 15 rounds at 2-, 3-, and 4-fold dilutions when EF-Tu encoding DNA was added (Fig. 7d, deep blue lines), whereas it continuously declined when GFP encoding DNA was used as a negative control (green lines). Furthermore, also in agreement with the prediction, translational activity at an 8-fold dilution declined even in the presence of EF-Tu encoding DNA. To further confirm the existence of a sustainable regeneration point, we performed the same experiment with a reduced initial EF-Tu concentration in round 1 from 80 µM to 0.625 μM and observed that translational activity increased toward the level observed in experiments that started from a higher concentration (Fig. 7d, light blue lines), consistent with convergence toward the predicted steady-state concentration. We then plotted the final translation activities at rounds 8–15 as the sustainable regeneration points in Fig. 7b by converting them to the corresponding EF-Tu concentration. The data points scattered around the model predictions within a 10-fold difference, supporting the reliability of our theoretical framework for predicting both regeneration feasibility and steady-state translation activity to a certain extent.

Additionally, to directly confirm the sustainable synthesis of EF-Tu during serial dilution, we analyzed samples from the 3-fold dilution condition by SDS-PAGE followed by CBB staining. A clear band was observed at the expected molecular weight of EF-Tu across all 15 rounds in samples containing EF-Tu encoding DNA (Fig. 7e). Quantification of the band intensities supported the constant EF-Tu existence (Fig. S8).

### Simultaneous regeneration experiment of EF-Tu and IF3

We next tested the applicability of our regeneration framework to the co-regeneration of two TFs, EF-Tu and IF3. However, a key challenge needed to be addressed: the expression curve, a central concept of the theoretical framework, is not directly applicable to the simultaneous regeneration of multiple TFs because the expression curve of each regenerating TF dynamically changes depending on the concentration of the other regenerating TFs. We addressed this challenge by ensuring that IF3 was always expressed in excess, so that the expression curve of EF-Tu remained independent of IF3 concentration. Under this condition, only the expression curve of EF-Tu needed to be considered for the feasibility of sustainable regeneration. This condition is readily achievable because EF-Tu is required at a substantially higher concentration than the other TFs in the PURE system (Fig. 2a).

We then determined the conditions under which IF3 would always be expressed in excess. Based on the dependence of translational activity on IF3 concentration (Fig. 4a) and the relative activity per protein of IF3 (1.6, Fig. 3e), we estimated that 0.5 µM of IF3 would be sufficient for the reaction. To express this amount of IF3, we selected 3.5 nM for the EF-Tu encoding DNA and 0.5 nM for the IF3 encoding DNA, which were predicted to express 6.1 µM of EF-Tu and 1.1 µM of IF3 under co-expression conditions, based on the co-expression data (Fig. 4c). Using these conditions, we derived the expression curve of EF-Tu at 2- and 3-fold dilutions to assess the feasibility of co-regeneration (Figs. 8a and S9). The predicted steady-state EF-Tu concentrations as a function of dilution rate are shown in Fig. 8b, predicting that sustainable co-regeneration of EF-Tu and IF3 would be possible over a dilution rate range of 1.8–5.1 fold.

**Figure 8.**
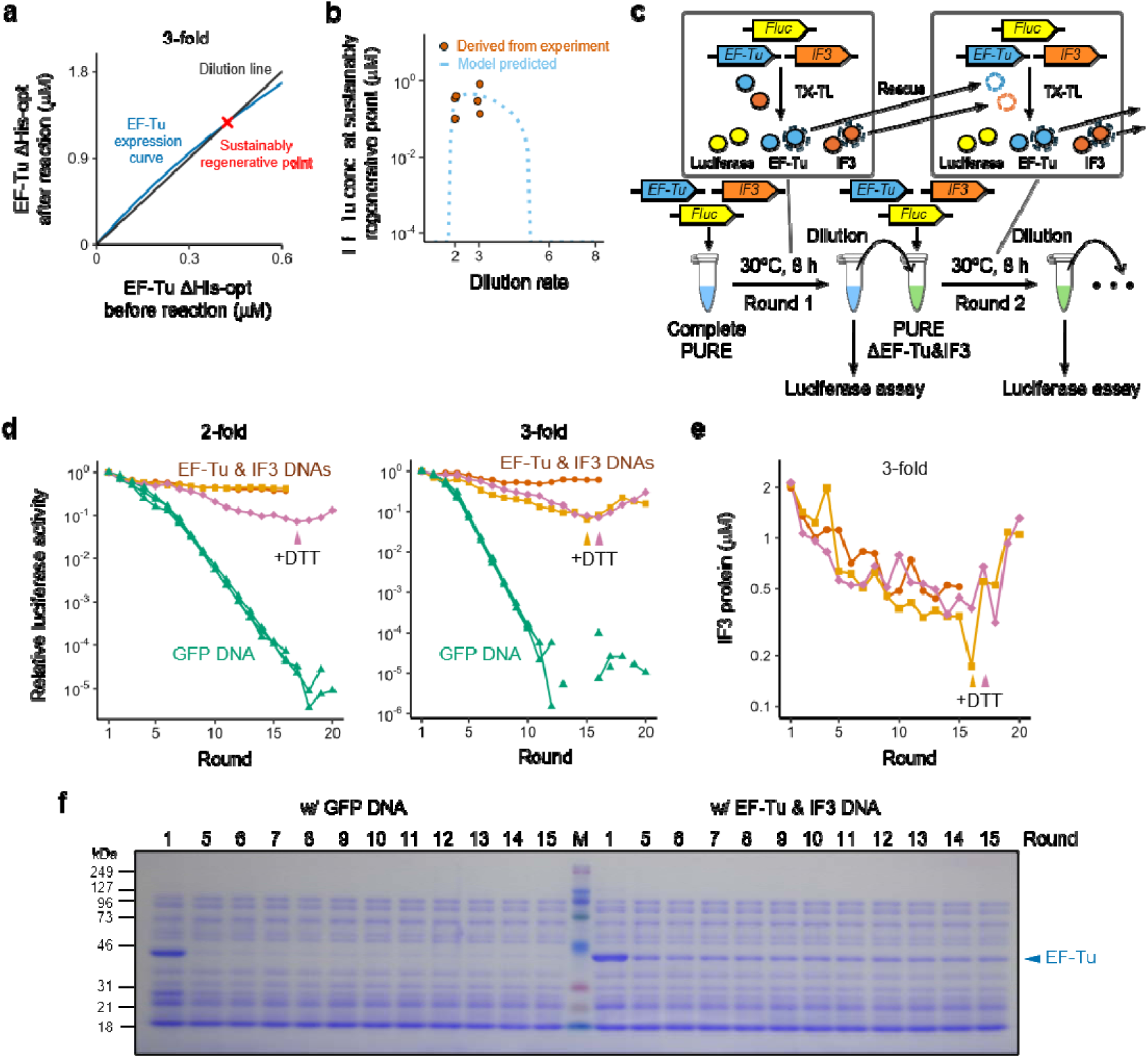
Prediction and experimental verification of sustainable co-regeneration of EF-Tu and IF3. (a) Theoretical prediction of co-regeneration of EF-Tu and IF3 at 3-fold dilution. The black line represents the dilution line (*y = 3x*), and the blue curve represents the EF-Tu expression curve at 3-fold dilution. The intersection point is indicated by a red cross. (b) Theoretical predictions and experimentally estimated steady-state EF-Tu concentrations during serial dilution. The dashed line represents the predicted EF-Tu concentration at the steady state. Orange circles indicate EF-Tu concentrations derived from the experimentally measured luciferase activities at the end of the serial dilution experiment in Fig. 8d. (c) Schematic of the co-regeneration serial dilution experiment. In the first round, EF-Tu encoding DNA (3.5 nM), IF3 encoding DNA (0.5 nM), and Fluc encoding DNA (0.01 nM) were expressed in the complete PURE system. In subsequent rounds, the reaction mixtures were diluted at 2- and 3-fold with the PURE system lacking EF-Tu and IF3 (PURE ΔEF-Tu&IF3) and containing the same DNA sets. All reactions were performed at 30°C for 8 h. Luciferase activity was measured after each round to assess translational activity. (d) Trajectories of luciferase activity at 2- and 3-fold dilution. Luciferase activities were normalized to the round 1 value. (e) Verification of IF3 synthesis. Samples from the 3-fold dilution experiment were analyzed by the HiBiT assay to quantify IF3 expression levels. Arrowheads indicate the rounds at which DTT supplementation (1 mM) was started. (f) Verification of EF-Tu synthesis. Samples from the 3-fold dilution experiment were subjected to SDS-PAGE on a 9% gel and stained with CBB. The band corresponding to EF-Tu is indicated by an arrowhead. Band intensities were quantified in Fig. S10.

To experimentally verify these predictions, we performed serial dilution experiments for up to 16–20 rounds at 2- and 3-fold dilutions (Fig. 8c). We used EF-Tu encoding DNA (3.5 nM), IF3 encoding DNA (0.5 nM), and luciferase encoding DNA (0.01 nM). From round 2 onwards, the reactions were diluted with a ΔEF-Tu&IF3 PURE system. Consistent with the theoretical framework predictions, translational activity was maintained for up to 16 rounds at both dilutions (Fig. 8d). In one or two independent experiments, we unexpectedly observed a gradual decline in activity by round 15, and the addition of 1 mM DTT restored the activity in these experiments (indicated as “+DTT”), suggesting that the decline was attributable in part to an oxidative environment accumulating through serial dilution. The final translational activity values were also plotted in Fig. 8b after converging to the corresponding EF-Tu concentrations, indicating that the values varied around the predicted curve within a 10-fold range. These demonstrate that our theoretical framework is extendable to the simultaneous regeneration of the two major TFs, EF-Tu and IF3. Direct detection of IF3 and EF-Tu was also conducted using a HiBiT tag attached to IF3 (Fig. 8e) and SDS-PAGE with CBB staining (Fig. 8f; quantification of band intensities is shown in Fig. S10), respectively. These data demonstrate that EF-Tu and IF3 were sustainably co-regenerated during the serial dilution process, consistent with the prediction.

## Discussion

Constructing a self-regenerating system is a significant challenge in synthetic biology. The lack of a theoretical framework capable of predicting the feasibility of regeneration from the characteristics of each component has been a major obstacle to expanding the number of regenerating proteins. In this study, we established such a theoretical framework for identifying the conditions required for sustainable regeneration of TFs. In this framework, the feasibility of regeneration is predicted from the intersection between the TF expression curve and the dilution line (Fig. 1b); and the same framework can moreover be extended to the simultaneous regeneration of multiple TFs by maintaining one TF as the limiting factor for translational activity. As experimental verification, we derived the expression curve from independently measured parameters, predicted the dilution rates permitting sustainable regeneration, and experimentally demonstrated sustainable regeneration of EF-Tu, the most abundant TF in the PURE system, for up to 15 rounds of serial dilution (Fig. 7) and sustainable co-regeneration of EF-Tu and IF3, the second most abundant essential TF in the PURE system, for up to 15–20 rounds (Fig. 8d). These results demonstrate that the proposed framework provides a rational and systematic methodology for the sustainable regeneration of major TFs, contributing to the development of self-regenerating artificial systems.

We found that increasing the concentration of T7 RNAP gradually decreased translational activity during serial dilution (Fig. 5b), and this decrease was more pronounced at lower dilution rates (Figs. 5c). One possible explanation is the depletion of substrates (e.g., NTP) by excess RNAP; however, substrate depletion alone is unlikely to explain this degree of decrease because the newly synthesized luciferase activity (i.e., translation activity) decreased by less than half at 2-fold dilution (Fig. 5d). A more likely explanation is the accumulation of inhibitory factors. One candidate inhibitory factor is untranslated mRNA, which may reduce translation efficiency by sequestering ribosomes and TFs^42^. If this is the case, introducing slower T7 RNAP mutants or an mRNA degradation system could alleviate the translation decline by suppressing the excessive accumulation of mRNA^27^. Another candidate factor is inorganic phosphate, which can sequester the free magnesium ions required for efficient translation^43^. In addition, a lowered pH in the reactions may also contribute to the decline in translational activity^19^. One approach to mitigate these effects is the use of a dialysis system, which would allow inorganic phosphate and hydrolase ions to diffuse into an external solution^39,44,45^. Alleviating this translational inhibition during serial dilution is therefore the next important challenge toward increasing the number of sustainably regeneratable TFs.

To achieve whole self-regeneration in the future, all TFs must be sustainably regenerated. The theoretical framework developed in this study enables a more realistic assessment of the requirements for achieving this goal. According to our theoretical framework and the measured parameters, sustainable regeneration of all non-ribosomal TFs is possible if two requirements are satisfied: (1) EF-Tu remains the rate-limiting factor for translation (i.e., all other TFs are sufficiently expressed) and (2) the expression curve of EF-Tu intersects a dilution line. The non-ribosomal TFs other than EF-Tu are included in the PURE system at 1 mg/mL in total (Table S3), and we assumed that this is a sufficient expression level. To satisfy condition (1), TFs other than EF-Tu must be expressed at levels exceeding 1 mg/mL. Therefore, to satisfy the condition (2), EF-Tu must be expressed more than approximately 0.2 mg/ml. However, using a batch reaction of the current PURE system, the maximum protein yield during serial dilution is approximately 0.1 mg/mL, which is much less than the required level. Therefore, a ∼14-fold increase would be required; our calculations indicate that at this level, the expression curve of EF-Tu would intersect the 3-fold dilution line (Fig. S11).

We consider this 14-fold increase to be an achievable goal, as several avenues for improvement exist. First, as described above, the steady-state translational activity was approximately halved relative to the initial level under 3-fold dilution conditions; resolving the inhibitory effects discussed above could therefore yield a 2-fold increase. Second, proteins synthesized in the PURE system often exhibit higher activity per protein compared to their purified counterparts (Fig. 3e). In addition to the His-tag removal and codon optimization performed in this study, the use of proteins from alternative bacteria^38^ and high-activity variants generated by in vitro evolution^46^ may further improve functional activity, thereby reducing the required fold-improvement in expression. Third, a dialysis system has been shown to increase expression levels approximately 3–10-fold increase^39,44,45^. Furthermore, other strategies to enhance expression level have been reported, including the supplementation of additional proteins^43,47–49^, and the introduction of an mRNA degradation system^42^. As an alternative approach, Ganesh and Maerkl reported that a diluted PURE system containing a minimum concentration of TFs can theoretically synthesize all non-ribosomal TFs^40^, which may reduce the expression level required for regeneration. By combining these improvements, we believe that the simultaneous sustainable regeneration of all non-ribosomal TFs is a realistic and achievable goal.

## Methods

### DNA preparation

The whole sequences of the plasmids used in this study are shown in Supplemental Data. Plasmids encoding EF-Tu (pET-EFTu) and IF3 (pET-IF3) were constructed by exchanging the vector region of the original plasmids used for PURE system preparation^18^ with the pET vector. The plasmid encoding EF-TuΔHis (pET-EFTuΔHis) and IF3ΔHis (pET-IF3ΔHis) were constructed by removing the histidine tag by PCR, followed by In-Fusion cloning (Takara). Plasmids encoding codon-optimized EF-Tu and IF3 (pET-EF-TuΔHis-opt and pET-IF3ΔHis-opt) were constructed by optimizing the ORF sequences for *E. coli* codon usage using CodHonEditor^41^, followed by gene synthesis and cloning into a pET vector (Twist Bioscience). The plasmid encoding mCherry (pET-mCherry) was synthesized using an artificial gene synthesis service (Twist Bioscience).

DNA templates were prepared by PCR amplification from the corresponding plasmids and quantified based on A_260_ using a NanoDrop spectrophotometer (Thermo Fisher Scientific). Unless otherwise noted, PCR products were purified using the FastGene Gel/PCR Extraction Kit (Nippon Genetics). DNA templates encoding GFP and mCherry were amplified from plasmids (pET-GFP^50^ and pET-mCherry, respectively) using primers 1 and 2. For experiments shown in Fig. 2b, the GFP template was amplified using primers 3 and 4, and the PCR product was purified using the QIAquick PCR Purification Kit (QIAGEN). For experiments using the ΔRF1 or ΔRF2 PURE system, GFP templates were amplified using primers 3 and 5 or primers 3 and 6, respectively. Luciferase DNA template was prepared from previously constructed plasmid^51^ using primers 3 and 7. DNA templates encoding EF-Tu and IF3 were amplified from the corresponding plasmids described above using primers 1 and 2. DNA templates with a C-terminal HiBiT tag were prepared by overlap extension PCR. The HiBiT fragment was amplified using primers 8 and 9, and DNA templates containing sequences complementary to the HiBiT fragment were amplified using primer 1 and gene-specific reverse primers (primer 10; IF3+His and IF3ΔHis, 11; IF3ΔHis-opt, 12; EF-Tu+His, 13; EF-TuΔHis, 14; IF3ΔHis-opt). The DNA templates were purified and fused with the HiBiT fragment by overlap extension PCR using primers 1 and 15.

### PURE system preparation

All components of the PURE system used in this study were individually purified by affinity chromatography with a histidine tag, followed by gel filtration, as previously described^38^. The compositions of the PURE system used in Fig. 2b (optimized for DNA replication coupling) are provided in Table S2, and those used in subsequent experiments (optimized for gene expression) are provided in Table S3. The total tRNA mixture (E. coli MRE 600) was purchased from Roche.

### Activity assay of EF-Tu and IF3

EF-Tu or IF3 DNA (4 nM) was incubated in the PURE system at 30°C for 8 h (1^st^ reaction). For EF-Tu expression, the EF-Tu concentration was reduced to 10 μM. mCherry DNA was used as a negative control. The reaction mixtures were then serially diluted (EF-Tu; 2-, 3-, and 4-fold, IF3; 6-, 8-, and 10-fold) using a protein dilution buffer (50 mM HEPES-KOH (pH 7.6), 100 mM KCl, 10 mM MgCl_2_, 7 mM 2-mercaptoethanol, 10 mg/mL BSA, 1 mM DTT, and 30% (v/v) glycerol). The diluted mixtures were added to PURE systems lacking the respective TFs but containing GFP encoding DNA (3 nM) and incubated at 30°C for 8 h (2^nd^ reaction). GFP fluorescence was measured every 10 min (Mx3005P, Agilent Technologies). GFP synthesis rates were determined from the slope of fluorescence intensity during the linear phase (EF-Tu; 3-5 h, IF3; 2-3 h). These rates were plotted against the dilution ratio, and the slope was defined as TF activity. For TFs synthesized in the PURE system, the expression levels were measured by parallel reactions using HiBiT-tagged templates under identical conditions. For purified proteins, 10 μM EF-Tu and 4.9 μM IF3 were used. Activity per protein was defined as the TF activity after subtraction of the negative control value and normalization to the corresponding expression level.

### Protein quantification via HiBiT assay

HiBiT-tagged TFs were quantified using the Nano-Glo HiBiT Lytic Detection System (Promega). The reaction mixture, SDS sample buffer (150 mM Tris-HCl (pH 7.4), 6% (w/v) SDS, 2.5 M 2-mercaptoethanol, 30% (v/v) glycerol, and a trace amount of bromophenol blue), and ultrapure water (Milli-Q water) were mixed in a 1:1:1 (v/v/v) ratio and incubated at 95°C for 5 min. The mixture was then diluted with BSA stock buffer (50 mM HEPES-KOH (pH 7.6), 100 mM KCl, 10 mM MgCl_2_, 7 mM 2-mercaptoethanol, 10 mg/mL BSA, and 30% (v/v) glycerol) and added to 20 μL of Nano-Glo HiBiT Lytic Reagent. Luminescence was measured using a GloMax luminometer (Promega). The protein concentration was calculated from the luminescence using a control HiBiT-tagged protein, the PgmA protein of E. coli. The control protein was expressed in E. coli and purified using a histidine tag by the same method as the PURE system component, but omitted the gel-filtration column. The protein concentration was estimated by A_280_. For IF3, we utilized HiBiT-tagged templates in all experiments intended for monitoring expression levels during serial dilution to confirm IF3 expression. This approach was adopted due to the overlap in molecular weight between IF3 (21 kDa) and other PURE components (ribosome recycling factor, 21 kDa; myokinase, 22 kDa), which prevents its clear identification by SDS-PAGE analysis.

### Parameter fitting

The data obtained in Figs. 4a and 4b were fitted to Michaelis–Menten-type saturation equations by nonlinear least-squares regression using the nls function in R version 4.2.1. The following equations were used:

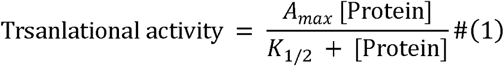

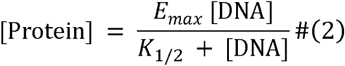

Here, *A_max_* and *E_max_* represent the maximum translational activity and maximum expression level, respectively. *K_1/2_*represents the protein or DNA concentration yielding a half-value of *A_max_* or *E_max_*.

### Co-expression model of EF-Tu and IF3

We assumed that competition among DNA templates for limited expression resources follows a competitive inhibition model. Under this assumption, the expression levels of each protein can be described by Eqs. 3 and 4, where parapeters *K_1/2_* and *E_max_* were obtained in Fig. 4b.

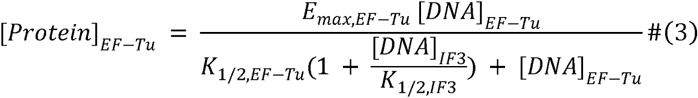

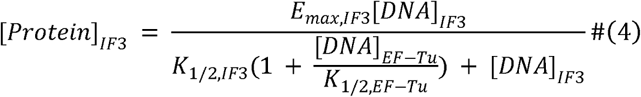

### Calculation of newly synthesized luciferase activity

To estimate newly synthesized luciferase activity, the residual luciferase activity after heat inactivation during 8 h incubation was first determined as follows. EF-Tu encoding DNA (4 nM) and Fluc encoding DNA (0.01 nM) were added to the PURE system and incubated at 30°C for 8 h, after which luciferase activity was measured (round 1). The reaction mixture was then further incubated at 30°C for 8 h, and luciferase activity was measured again (round 2). The ratio of luciferase activity in round 2 to that in round 1 was defined as the residual activity, which was 0.81 ± 0.02 (mean ± SD, n = 3). Because the luciferase protein derived from the previous round was carried over into the next round at 1/D (D: Dilution rate), newly synthesized luciferase activity in round N can be described by Eq. 5.

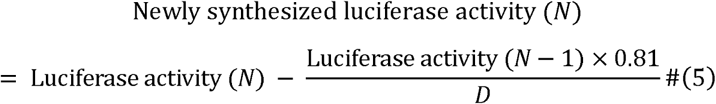

### Estimation of the steady-state translational activity during serial dilution

Assuming that the amount of inhibitor, *I*, accumulates in each round, the inhibitor concentration before the reaction in round *N* (*I_N_*) can be described by Eq. 6.

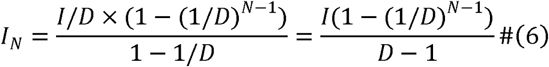

Assuming exponential inhibition of translational activity by the accumulated inhibitor, the translational activity in round *N* can be described by Eq. 7, where *c* is an inhibition coefficient. Because only the product *cI* appears in the equation, it was treated as a single fitting parameter, *k*.

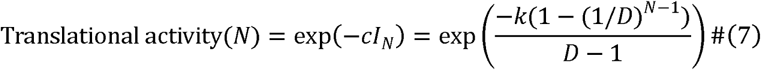

### Serial dilution experiments

In round 1, TF-DNA (4 nM) and Fluc-DNA (0.01 nM) were added to the Complete PURE system and incubated at 30°C for 8 h. GFP-DNA was used as a negative control. From round 2 onwards, the reaction mixture from the previous round was transferred into Complete PURE (Figs. 5b and 5c) or ΔPURE lacking the target TFs (Figs. 7d and 8d), both of them containing template DNAs, at a defined dilution rate and incubated at 30°C for 8 h. After each reaction, a 1 μL aliquot of the reaction mixture was added to 20 μL of luciferase assay reagent (Promega), and luminescence was measured using a GloMax luminometer (Promega).

### SDS-PAGE analysis

The reaction mixture, SDS sample buffer, and ultrapure water (Milli-Q water) were mixed in a 1:1:1 (v/v/v) ratio and incubated at 95°C for 5 min. A 2 μL aliquot of the mixture was subjected to SDS-polyacrylamide gel electrophoresis (SDS-PAGE) using 9% gel. Proteins were stained with Coomassie Brilliant Blue (CBB), and band intensities were analyzed using ImageJ.

## Supporting Information

Supporting Information includes supplemental figures (Figures S1 – S11) and tables (Tables S1 – S3).

## Author contribution

KS and NI planned the experiments and wrote the manuscript. KS and KH performed the experiments.

## Acknowledgement

We thank Ms. Kayo Aoyama and Ayu Saito for their technical support.

