## Supplemental Figures for "Theoretical framework and experimental demonstration of sustainable in vitro regeneration of major translation factors EF-Tu and IF3"

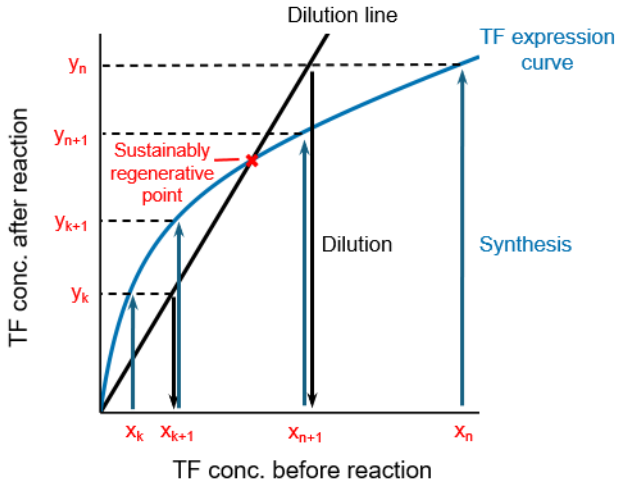


**Figure S1. Detailed schematic of the theoretical framework of TF regeneration.**

The blue curve represents the TF expression curve, showing the TF concentration after the reaction as a function of the TF concentration before the reaction. If the reaction begins at a given TF concentration at round n ($x_n$), the TF concentration after a single round of TX-TL reaction becomes the corresponding y-value on the expression curve ($y_n$). Following dilution at a given dilution rate (D), the initial TF concentration for the next round (n+1) becomes $x_{n+1}$, the x-coordinate of the intersection of $y = y_n$ and the dilution line, $y = Dx$. In the following round of TX-TL reaction, the TF concentration increases to $y_{n+1}$. Through this process, both x and y decrease and converge to the intersection point between the expression curve and the dilution line (the sustainably regenerative point). Conversely, if the reaction begins at a lower TF concentration ($x_k$), the TF concentration after a single round of TX-TL becomes $y_k$. Upon dilution, the concentration becomes $x_{k+1}$, which is greater than $x_k$, thereby driving the TF concentration toward the sustainably regenerative point. Therefore, if the expression curve intersects with a dilution line, the intersection point defines the steady-state TF concentration.

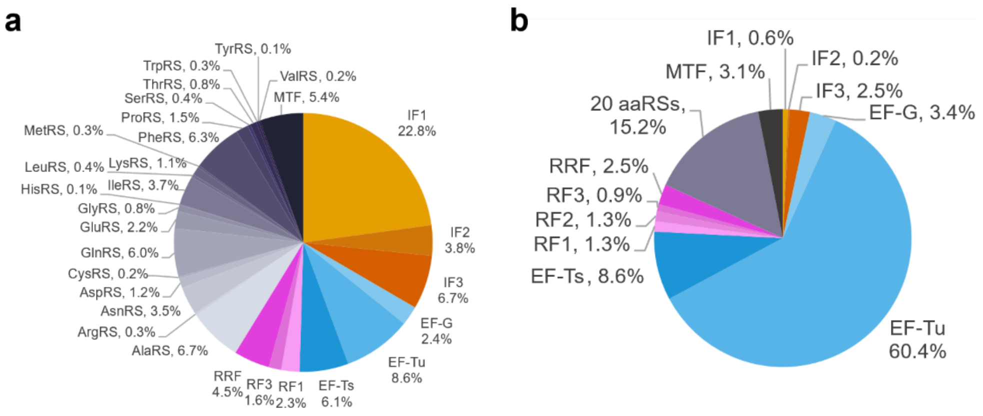


**Figure S2. TF compositions in other PURE systems.**

(a) The original TF composition reported by Shimizu et al.[1] (b) The alternative TF composition reported by Lavickova et al.[2] Both compositions are expressed as molar concentrations. The TF composition of the PURE system used in this study is shown in Fig. 2a. IF, initiation factor; EF, elongation factor; RF, release factor; RRF, ribosome recycling factor; aaRSs, aminoacyl-tRNA synthetases; MTF, methionyl-tRNA formyltransferase.

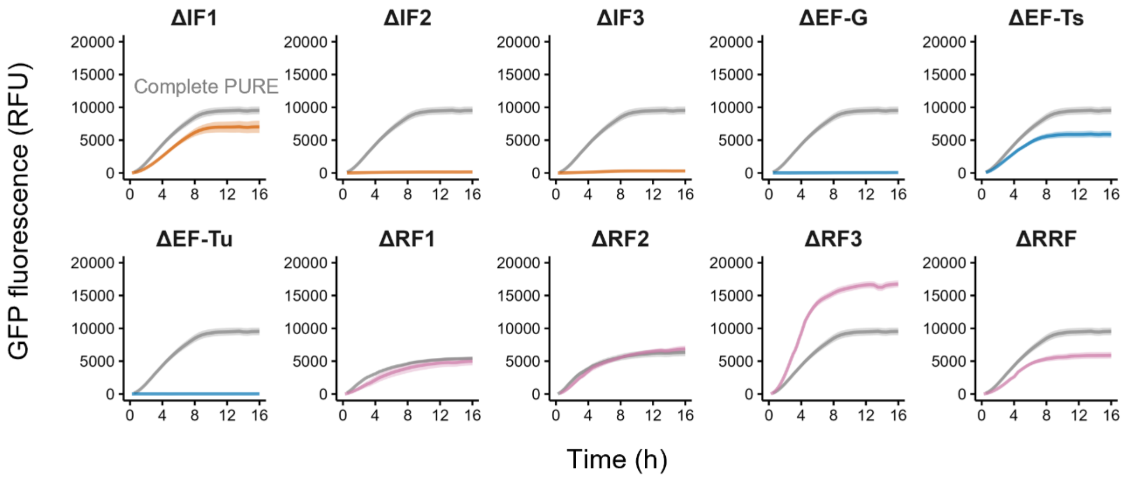


**Figure S3. Time−course data of GFP fluorescence in TF-depleted PURE systems.**

GFP encoding DNA (3 nM) was added to the PURE system lacking each individual TF (ΔTF) and incubated at 30°C for 16 h. For ΔRF1 and ΔRF2, GFP encoding DNAs containing stop codons specifically recognized by RF1 or RF2 were used. GFP fluorescence was measured every 15 min. The gray curves indicate GFP fluorescence in the complete PURE system containing all TFs (Complete PURE). Data points represent mean values ± SD from three independent experiments.

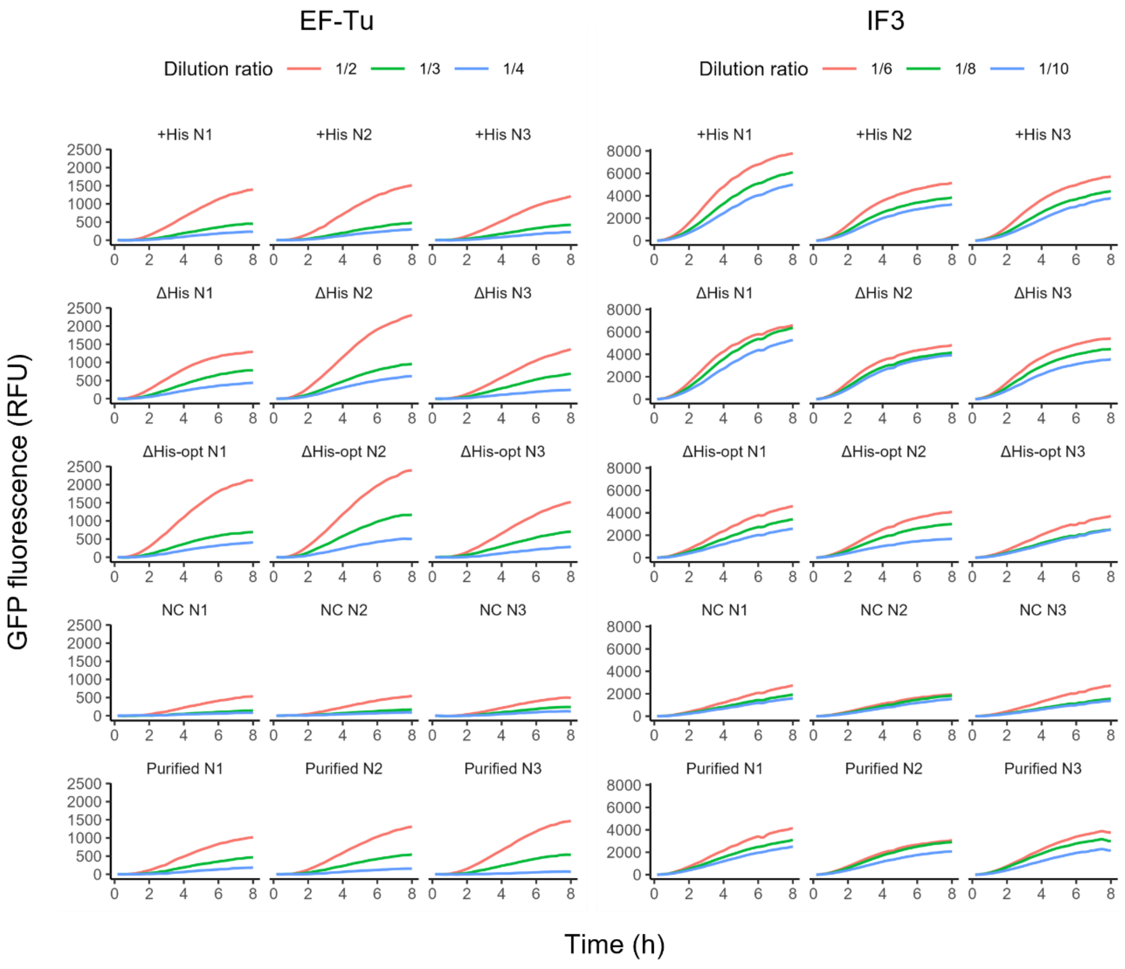


**Figure S4. Time-course data of GFP fluorescence during the 2nd reaction in the activity assay.**

Dilution series of the 1st reaction mixtures were added to the PURE systems lacking the corresponding target TF and containing GFP encoding DNA. The reaction mixtures were incubated at 30°C for 8 h. GFP fluorescence was measured every 10 min.

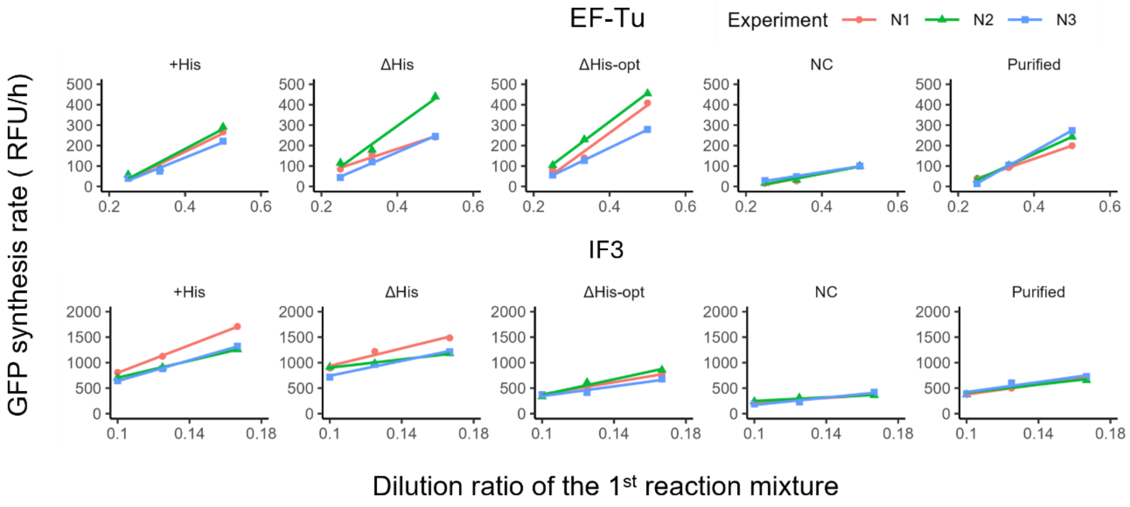


**Figure S5. GFP synthesis rates plotted against dilution ratio.**

GFP synthesis rates were calculated from the slopes of the linear regions (EF-Tu: 3–5 h, IF3: 2–3 h) of the GFP fluorescence time courses shown in Fig. S4. The slope obtained by linear regression of the GFP synthesis rates at the three dilution ratios was defined as the TF activity value reported in Fig. 3d.

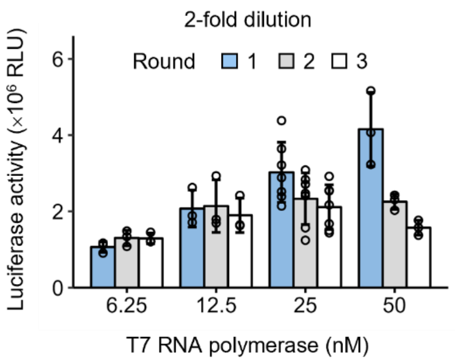


**Figure S6. Effect of T7 RNAP concentration on luciferase activity at 2-fold dilution.**

Luciferase activity was measured up to round 3 in reactions containing 6.25, 12.5, 25, or 50 nM T7 RNAP. Bars indicate mean values ± SD from three to seven independent experiments. The corresponding results under 3-fold dilution are shown in Fig. 5b.

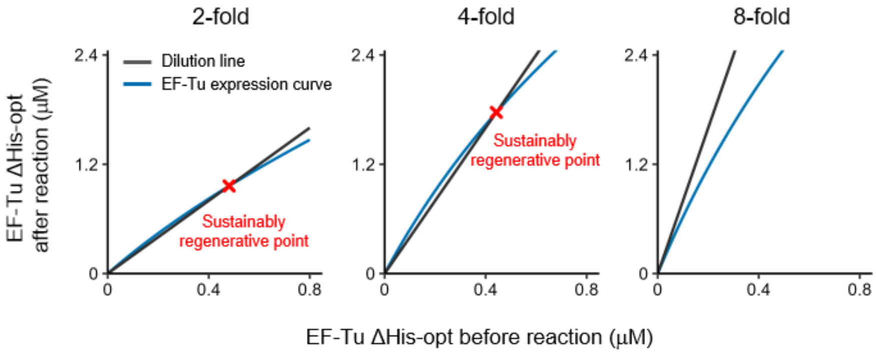


**Figure S7. Theoretical predictions of EF-Tu regeneration at 2-, 4-, and 8-fold dilutions.**

The black line represents the dilution line ($y = Dx$; $D = 2$, 4, or 8), and the blue curve represents the EF-Tu expression curve at each dilution rate, derived in Fig. 6e. Intersection points are indicated by a red cross. The theoretical prediction at 3-fold dilution is shown in Fig. 7a.

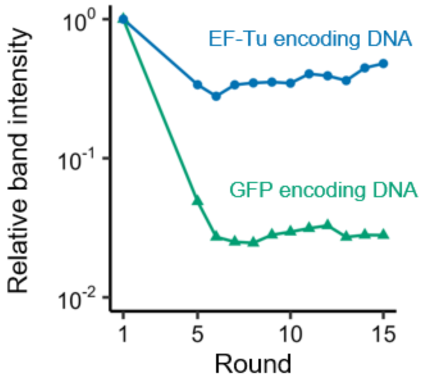


**Figure S8. Quantification of EF-Tu band intensities from Fig. 7e.**

The band intensities corresponding to EF-Tu were quantified using ImageJ and normalized to the round 1 value.

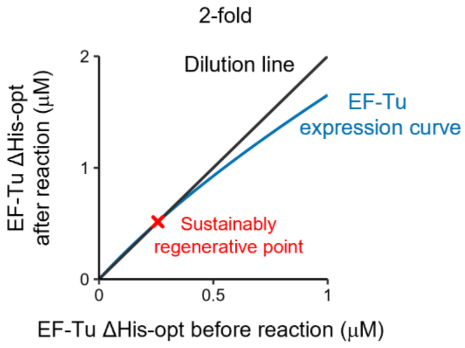


**Figure S9. Theoretical prediction of co-regeneration of EF-Tu and IF3 at 2-fold dilution.**

The black line represents the dilution line (y = 2x), and the blue curve represents the EF-Tu expression curve at 2-fold dilution. The intersection point is indicated by a red cross. The corresponding theoretical prediction at 3-fold dilution is shown in Fig. 8a.

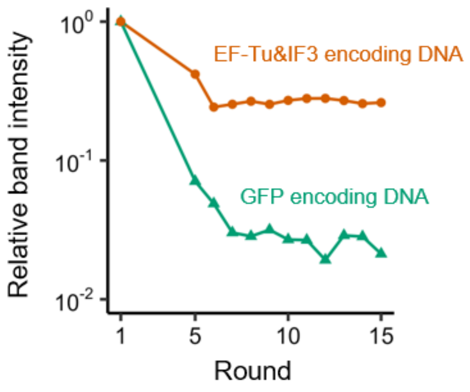


**Figure S10. Quantification of EF-Tu band intensities from Fig. 8f.**

The band intensities corresponding to EF-Tu were quantified using ImageJ and normalized to the round 1 value.

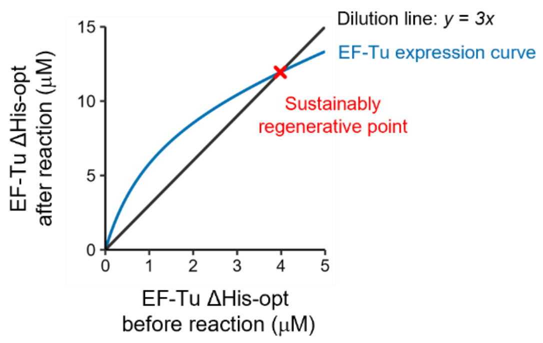


**Figure S11. Theoretical prediction of EF-Tu regeneration assuming a 14-fold increase in translational activity.**

The DNA concentrations were set to 1 nM for EF-Tu encoding DNA and 3 nM for TF encoding DNAs other than EF-Tu. The black line represents the dilution line (y = 3x), and the blue curve represents the EF-Tu expression curve assuming a 14-fold increase in translational activity. The intersection point is indicated by a red cross.

**Table S1. Primer sets used in this study.**

| Number | Sequence |
|---|---|
| 1 | GCGTCCGGCGTAGAGGATC |
| 2 | TCCGGATATAGTTCCTCCTTTCAG |
| 3 | GCGAAATTAATACGACTCACTATAGGG |
| 4 | GGTTATGCTAGTTATTGCTCAGCGG |
| 5 | GAGCTCGAATTCACTATTTGTAGAGCTC |
| 6 | GAGCTCGAATTCATCATTTGTAGAGCTC |
| 7 | GAGCAGACAAGCCCGTCAG |
| 8 | GGTTCTGGTGGTAATTCTGGTTCTTCTGGTGGCTCCTCTGGTGTTTCTGGTTGGC |
| 9 | GGCCGCAAGCCTATTAAGAAATTTTCTTAAATAAACGCCAACCAGAAACACCAGAG |
| 10 | GAACCAGAATTACCACCAGAACCCTGTTTCTTCTTAGGAGCGAGC |
| 11 | GAACCAGAATTACCACCAGAACCCTGTTTTTTTTCGGAGCCAGAAC |
| 12 | GAACCAGAATTACCACCAGAACCGTGATGGTGATGAGATCTGCCC |
| 13 | GAACCAGAATTACCACCAGAACCGCCCAGAACTTTTGCTACAACG |
| 14 | GAACCAGAATTACCACCAGAACCACCCAGAACTTTAGCTACAACGCC |
| 15 | GGCCGCAAGCCTATTAAGAA |

**Table S2. Composition of the customized PURE system used in Fig. 2b**

| Component | Concentration | Component | Concentration |
|---|---|---|---|
| Initiation factor 1 | 25 µM (200 µg/mL) | Tyrosyl-tRNA synthetase | 0.15 µM (7.2 µg/mL) |
| Initiation factor 2 | 1.0 µM (100 µg/mL) | Valyl-tRNA synthetase | 17 nM (1.9 µg/mL) |
| Initiation factor 3 | 4.9 µM (100 µg/mL) | Methionyl-tRNA Formyltransferase | 0.59 µM (20 µg/mL) |
| Elongation factor G | 1.1 µM (83 µg/mL) | Myokinase | 1.4 µM (30 µg/mL) |
| Elongation factor Tu | 80 µM (3500 µg/mL) | Creatine kinase | 0.25 µM (11 µg/mL) |
| Elongation factor Ts | 3.3 µM (100 µg/mL) | Nucleoside diphosphate kinase | 16 nM (0.2 µg/mL) |
| Release factor 1 | 49 nM (2.0 µg/mL) | Pyrophosphatase | 41 nM (1.3 µg/mL) |
| Release factor 2 | 48 nM (2.0 µg/mL) | Trigger Factor | 1.0 µM (48 µg/mL) |
| Release factor 3 | 0.17 µM (10 µg/mL) | E. coli DEAH type RNA helicase A | 63 nM (9.2 µg/mL) |
| Ribosome recycling factor | 3.9 µM (80 µg/mL) | T7 RNA polymerase | 0.42 U/µL |
| Alanyl-tRNA synthetase | 0.73 µM (70 µg/mL) | Ribosome | 1.0 µM |
| Arginyl-tRNA synthetase | 31 nM (2.0 µg/mL) | Tyrosine and cysteine | 0.3 mM each |
| Asparaginyl-tRNA synthetase | 0.42 µM (22 µg/mL) | 18 other amino acids | 0.36 mM each |
| Asparagyl-tRNA synthetase | 0.12 µM (8.0 µg/mL) | tRNA mix (Roche, *E. coli*) | 0.52 mg/mL |
| Cysteinyl-tRNA synthetase | 24 nM (1.3 µg/mL) | ATP | 0.375 mM |
| Glutaminyl-tRNA synthetase | 60 nM (3.8 µg/mL) | GTP | 0.25 mM |
| Glutamyl-tRNA synthetase | 0.23 µM (13 µg/mL) | CTP | 0.125 mM |
| Glycyl-tRNA synthetase | 86 nM (9.6 µg/mL) | UTP | 0.125 mM |
| Histidyl-tRNA synthetase | 85 nM (4.0 µg/mL) | N-2-hydroxyethylpiperazineN'-2-ethanesulfonic acid (pH 7.6) | 100 mM |
| Isoleucyl-tRNA synthetase | 0.37 µM (38 µg/mL) | Potassium glutamate | 70 mM |
| Leucyl-tRNA synthetase | 41 nM (4.0 µg/mL) | Spermidine | 0.375 mM |
| Lysyl-tRNA synthetase | 0.12 µM (6.6 µg/mL) | Magnesium acetate | 10.8 mM |
| Methionyl-tRNA synthetase | 0.11 µM (8.3 µg/mL) | Creatine phosphate | 25 mM |
| Phenylalanyl-tRNA synthetase | 0.13 µM (17 µg/mL) | Dithiothreitol | 6 mM |
| Prolyl-tRNA synthetase | 0.13 µM (11 µg/mL) | 10-formyl-5,6,7,8-tetrahydro folic acid | 10 µg/mL |
| Seryl-tRNA synthetase | 78 nM (3.8 µg/mL) | Yeast inorganic pyrophosphatase (NEB) | 0.2 mU/µL |
| Threonyl-tRNA synthetase | 84 nM (6.2 µg/mL) | RNase inhibitor (Promega) | 0.1 U/µL |
| Tryptophanyl-tRNA synthetase | 28 nM (1.0 µg/mL) | EDTA | 2 mM |

**Table S3. Composition of the customized PURE system used other than Fig. 2b**

| Component | Concentration | Component | Concentration |
|---|---|---|---|
| Initiation factor 1 | 25 µM (200 µg/mL) | Tyrosyl-tRNA synthetase | 0.15 µM (7.2 µg/mL) |
| Initiation factor 2 | 1.0 µM (100 µg/mL) | Valyl-tRNA synthetase | 17 nM (1.9 µg/mL) |
| Initiation factor 3 | 4.9 µM (100 µg/mL) | Methionyl-tRNA Formyltransferase | 0.59 µM (20 µg/mL) |
| Elongation factor G | 1.1 µM (83 µg/mL) | Myokinase | 1.4 µM (30 µg/mL) |
| Elongation factor Tu | 80 µM (3500 µg/mL) | Creatine kinase | 0.25 µM (11 µg/mL) |
| Elongation factor Ts | 3.3 µM (100 µg/mL) | Nucleoside diphosphate kinase | 16 nM (0.2 µg/mL) |
| Release factor 1 | 49 nM (2.0 µg/mL) | Pyrophosphatase | 41 nM (1.3 µg/mL) |
| Release factor 2 | 48 nM (2.0 µg/mL) | Trigger Factor | 1.0 µM (48 µg/mL) |
| Release factor 3 | 0.17 µM (10 µg/mL) | E. coli DEAH type RNA helicase A | – |
| Ribosome recycling factor | 3.9 µM (80 µg/mL) | T7 RNA polymerase | 25 nM (2.5 µg/mL) |
| Alanyl-tRNA synthetase | 0.73 µM (70 µg/mL) | Ribosome | 1.0 µM |
| Arginyl-tRNA synthetase | 31 nM (2.0 µg/mL) | Tyrosine and cysteine | 0.3 mM each |
| Asparaginyl-tRNA synthetase | 0.42 µM (22 µg/mL) | 18 other amino acids | 0.36 mM each |
| Asparagyl-tRNA synthetase | 0.12 µM (8.0 µg/mL) | tRNA mix (Roche, *E. coli*) | 3.12 mg/mL |
| Cysteinyl-tRNA synthetase | 24 nM (1.3 µg/mL) | ATP | 3.75 mM |
| Glutaminyl-tRNA synthetase | 60 nM (3.8 µg/mL) | GTP | 2.5 mM |
| Glutamyl-tRNA synthetase | 0.23 µM (13 µg/mL) | CTP | 1.25 mM |
| Glycyl-tRNA synthetase | 86 nM (9.6 µg/mL) | UTP | 1.25 mM |
| Histidyl-tRNA synthetase | 85 nM (4.0 µg/mL) | N-2-hydroxyethylpiperazineN'-2-ethanesulfonic acid (pH 7.6) | 100 mM |
| Isoleucyl-tRNA synthetase | 0.37 µM (38 µg/mL) | Potassium glutamate | 280 mM |
| Leucyl-tRNA synthetase | 41 nM (4.0 µg/mL) | Spermidine | 1.5 mM |
| Lysyl-tRNA synthetase | 0.12 µM (6.6 µg/mL) | Magnesium acetate | 18 mM |
| Methionyl-tRNA synthetase | 0.11 µM (8.3 µg/mL) | Creatine phosphate | 25 mM |
| Phenylalanyl-tRNA synthetase | 0.13 µM (17 µg/mL) | Dithiothreitol | 1.5 mM |
| Prolyl-tRNA synthetase | 0.13 µM (11 µg/mL) | 10-formyl-5,6,7,8-tetrahydro folic acid | 10 µg/mL |
| Seryl-tRNA synthetase | 78 nM (3.8 µg/mL) | Yeast inorganic pyrophosphatase (NEB) | – |
| Threonyl-tRNA synthetase | 84 nM (6.2 µg/mL) | RNase inhibitor (Promega) | – |
| Tryptophanyl-tRNA synthetase | 28 nM (1.0 µg/mL) | EDTA | 2 mM |
